# Sequential VEGF-A165 plasmid and AAV-follistatin gene therapy enhances muscle hypertrophy and capillarisation in C57BL/6 mice

**DOI:** 10.64898/2026.08.26.747237

**Authors:** Anna Vakhrusheva, Andrey A. Nedorubov, Vladimir Leshko, Ivan Morgunov

## Abstract

**Introduction:** Skeletal muscle loss in sarcopenia and neuromuscular disorders remains a major unmet medical need. AAV9-delivered follistatin (FST), a myostatin/activin antagonist, induces muscle hypertrophy; however, fibre growth without adequate vascular adaptation may limit therapeutic efficacy. We evaluated whether co-administration of a VEGF-A165 plasmid enhances the hypertrophic and angiogenic effects of intramuscular AAV-FST gene transfer in C57BL/6 mice.

**Methods:** Thirty-six C57BL/6 mice (18 males, 18 females) were assigned to PBS vehicle (n=10), AAV-FST (1×10^11^ vg; n=10), VEGF plasmid (100 μg; n=6), or combination treatment (VEGF plus AAV-FST; n=10). The contralateral hindlimb served as an internal control. Endpoints at Day 115 included hindlimb muscle mass ratio (R/L), transgene expression, FST protein levels, muscle fibre morphometry, capillary density, and safety assessments.

**Results:** Combination therapy produced the highest R/L ratio (1.176 ± 0.091; p=0.004; d=2.04), whereas AAV-FST alone showed a borderline effect (R/L=1.113; p=0.050). Compared with AAV-FST monotherapy, combination treatment increased muscle FST mRNA ~2.1-fold, protein levels ~2.0-fold, and muscle-to-liver expression ratio 2.6-fold. It also induced larger muscle fibres and doubled CD31^+^ vessel counts versus AAV-FST alone, indicating simultaneous hypertrophy and angiogenesis. No adverse haematological, biochemical, or histopathological findings were observed.

**Discussion:** Combined AAV-FST and VEGF therapy enhanced local muscle hypertrophy, increased capillary density, and improved the muscle-to-liver transgene expression profile compared with AAV-FST monotherapy. The regimen was well tolerated and supports further evaluation of angiogenic preconditioning as a strategy to improve muscle-directed gene therapy for muscle-wasting disorders.

## Introduction

Skeletal muscle wasting is a major clinical burden: sarcopenia affects 10–27% of adults aged ≥60 years and is associated with falls, frailty, and all-cause mortality (Cruz-Jentoft, 2019; Petermann-Rocha, 2021; Yuan, 2023; Stuck, 2023; Ethgen, 2016). No pharmacological agent is approved for sarcopenia (Rooks, 2019); the most advanced candidate, bimagrumab, increased lean mass but failed to improve physical function in phase II trials (Rooks, 2020; Rooks, 2017). The same myostatin/activin pathway is implicated in Duchenne/Becker muscular dystrophies (Cirsafulli, 2020; Salari, 2022) and cancer cachexia (Fearon, 2011; Huo, 2025), underscoring a broad unmet therapeutic need.

Follistatin is a high-affinity natural inhibitor of myostatin and the related ligand activin A; it acts both upstream — by sequestering circulating ligand from the activin type II receptor — and intracellularly, suppressing SMAD2/3 transcriptional activity and de-repressing the AKT/mTOR/p70S6K protein-synthesis axis (Lee, 2001; Trendelenburg, 2009; Sartori, 2009). Because follistatin neutralises both myostatin and activin, it produces a more pronounced hypertrophy than myostatin blockade alone in mdx and wild-type animals (Gilson, 2009; Lee, 2010; Iskenderian, 2018). Two follistatin isoforms — FST-288 (heparan-sulfate-binding, locally retained) and FST-315 (circulating) — have been used preclinically and clinically (Sidis, 2006; Datta-Mannan, 2013; Pearsall, 2019).

Early-phase human trials of AAV1-FS344 in Becker muscular dystrophy and inclusion-body myositis established an acceptable safety profile and provided proof-of-principle for muscle-directed AAV-follistatin therapy (Kota, 2009; Mendell, 2015; Al-Zaidy, 2015; Mendell, 2017; Greenberg, 2017). We selected AAV9 for its well-characterised skeletal muscle tropism, durable single-dose expression, and low pre-existing seroprevalence (Wang, 2019; Zinkarelli, 2008; Inagaki, 2006; Tabebordbar, 2021; Wang, 2024). AAV9 also serves as the vector backbone of the first systemically approved gene therapy, onasemnogene abeparvovec, supporting its translational relevance (Blair, 2022).

A potential limitation of purely anabolic strategies is that muscle fibre growth and vascular adaptation are not necessarily coupled. Experimental inhibition of the myostatin/activin pathway with soluble ActRIIB-Fc rapidly reduces capillary density and VEGF-A protein in mouse muscle (Hulmi, 2012). Likewise, myostatin-null muscle exhibits reduced specific force attributable in part to inadequate capillarisation (Matsakas, 2012). Capillary density and fibre size are co-regulated through exercise-induced VEGF signalling, suggesting that follistatin-driven hypertrophy may be limited by a rarefied capillary bed — a biological bottleneck for anabolic therapies (Olfert, 2010; Hoier, 2014).

Several studies support simultaneous targeting of anabolic and angiogenic pathways. Borselli et al. showed that co-delivery of VEGF and IGF-1 to ischaemic hindlimb outperformed either factor alone (Borselli, 2009), and Männistö et al. (2025) demonstrated that AAV-VEGF-B co-delivered with a myostatin-propeptide AAV rescued capillary rarefaction and enhanced muscle growth in healthy and diabetic mice (Männistö, 2025). Together, these findings suggest that coordinated stimulation of angiogenesis and muscle growth may provide advantages over activation of either pathway alone.

To stimulate angiogenesis, we selected pl-VEGF165 (Neovasculgen®), a CMV-driven plasmid encoding human VEGF-A165, registered in Russia in 2011 for peripheral arterial disease and supported by Phase IIb/III data (+110% pain-free walking at 6 months (Deev, 2015), a 210-patient post-marketing study (Deev, 2017), and 5-year follow-up with no tumour formation or aberrant angiogenesis (Deev, 2018). Plasmid-based delivery is self-limiting, contrasting with sustained AAV-VEGF overexpression, which potentially can cause haemangiomas (Satkauskas, 2001; Son, 2005; Olea, 2009; Karvinen, 2011; Gianni-Barrera, 2016).

To our knowledge, the combination of AAV-follistatin with a VEGF-A165 plasmid has not been reported. We hypothesised that sequential VEGF-induced angiogenic preconditioning followed by intramuscular AAV9-FST344 administration would enhance local muscle hypertrophy and increase muscle-targeted transgene expression compared with AAV-FST monotherapy. To test this, we characterised local skeletal muscle hypertrophy, microvascular density, FST transduction (mRNA and protein), and systemic safety after intramuscular AAV-FST (1×10^11^ vg) alone or in sequential combination with pl-VEGF165 (100 μg) in C57BL/6 mice of both sexes.

## Materials and Methods

### Test Articles

AAV-Follistatin (AAV-FST) was a recombinant adeno-associated virus serotype 9 (AAV9) encoding the human FST344 isoform, administered at a dose of 1×10^11^ vg per mouse in a total volume of 250 µL PBS, delivered across 5 intramuscular injection sites (50 µL per site). The VEGF plasmid Neovasculgen® (NexTgen, Russia) encodes VEGF165 and was administered as 100 µg in a total volume of 250 µL PBS (pH 7.4), likewise delivered across 5 intramuscular sites (50 µL per site). PBS (pH 7.4) served as the vehicle for all test articles; animals in Group 1 received PBS injections only and served as the vehicle control group.

### Animals

Thirty-six adult C57BL/6 mice (18 males, 18 females; 6–10 weeks; 20–24 g) were obtained from certified suppliers and housed under standard conditions (20–24°C, 45–65% relative humidity, 12 h light/dark, food and water ad libitum). All procedures were approved by the Institutional Animal Ethics Committee (Protocol No. 235, 24.10.2025) and complied with the Guide for the Care and Use of Laboratory Animals (8th edition, NRC, 2011) and GOST 33216-2014.

### Experimental design

Mice were randomly assigned to four groups (Table 1). Intramuscular injections were administered into the right hindlimb on the designated study days, with the contralateral hindlimb injected with PBS and used as an internal paired control. Body weight and clinical observations were recorded at baseline, after quarantine, at group allocation, and during the study (including Days 32 and 39). Haematology and serum biochemistry were evaluated on Days 32 (limited biochemistry panel), 55, and 115 (full panel). An interim necropsy was performed on Day 55, when 2 males and 2 females from each of Groups 1, 2, and 4 were euthanised for clinical pathology, organ weights, histopathology, and analysis of FST transgene and protein expression. Remaining animals from Groups 1, 2, and 4 and all animals from Group 3 underwent terminal assessment on Day 115, including full clinical pathology, organ weights, gross necropsy, histopathology, and collection of injected and contralateral hindlimb muscles for immunohistochemistry and muscle mass measurements. Group 3 (VEGF monotherapy) did not undergo interim necropsy.

**Table 1.** Experimental groups.

| Group | Treatment | Dose | Days of administration | n (M/F) |
| --- | --- | --- | --- | --- |
| 1 | Vehicle (PBS) | — | Days 1, 10, 25 | 10 (5M/5F) |
| 2 | AAV-FST monotherapy | $1 \times 10^{11}$ vg | Day 25 | 10 (5M/5F) |
| 3 | VEGF plasmid (Neovasculgen®) | 100 µg | Days 1, 10 | 6 (3M/3F) |
| 4 | Combo (VEGF + AAV-FST) | VEGF 100 µg + AAV-FST $1 \times 10^{11}$ vg | Days 1, 10 (VEGF); Day 25 (AAV-FST) | 10 (5M/5F) |

### Safety Assessments

#### Haematology and body weight

Blood was collected into EDTA tubes and analysed on a HemaLite 1280 haematology analyser. Measured parameters included WBC with differential (LYM, MID, GRAN), RBC, HGB, HCT, MCV, MCH, MCHC, RDW-SD, RDW-CV, PLT, MPV, PDW, PCT, and P-LCR. Time points were as described above. Body weight and clinical signs were recorded at each clinical pathology time point and at interim and terminal necropsy.

#### Serum biochemistry

The serum was separated by centrifugation and analysed on an ERBA XL-100 clinical chemistry analyser. In mice, a reduced panel (ALP, urea, creatinine, total bilirubin, ALT, AST, GGT) was used on Day 32, and a full panel (ALP, urea, creatinine, glucose, triglycerides, cholesterol, total bilirubin, total protein, albumin, ALT, AST, GGT, creatine kinase, Na^+^, K^+^, Cl^−^) on Days 55 and 115.

#### Histopathology

At necropsy, tissues (heart, lung, liver, kidney, brain, thymus, spleen, gonads, dorsal root ganglia, and injection-site skeletal muscle) were fixed in 10% neutral buffered formalin, processed, paraffin-embedded, sectioned at 3–4 µm, stained with haematoxylin–eosin, and examined by light microscopy. Mice were evaluated at the interim (Day 55) and terminal (Day 115) time points.

### Efficacy Assessments

#### Muscle mass

At terminal sacrifice, both hindlimbs were dissected, and the muscles of the lower hindlimb were excised and weighed to the nearest 0.1 mg using an analytical balance (Ohaus Scout SPX222). The right-to-left muscle mass ratio (R/L) was calculated as the primary efficacy endpoint. Additionally, the following organs were excised and weighed: heart, lungs, liver, kidneys, thymus, spleen, gonads, and brain. Paired organs were weighed together. Between-group comparisons of organ weights were performed using the Kruskal–Wallis test.

#### Immunohistochemistry

Cryosections (7–10 µm) of injected and contralateral hindlimb muscles were fixed in 4% paraformaldehyde for 10–15 min, permeabilised with 0.1% Triton X-100 in PBS for 5 min, blocked with 2% BSA/PBS for 30 min, and stained with a mixture of primary antibodies against laminin (PAA082Mu01, Cloud-Clone Corp., China) and CD31 (ab182981, Abcam, UK) at a dilution of 1:500. Secondary antibodies conjugated to Alexa Fluor 488 (laminin) (ab150077, Abcam, UK) and Alexa Fluor 594 (CD31) (ab150116, Abcam, UK) were applied for 45 min, followed by DAPI (Lumiprobe, Russia) counterstaining. Fibre size was quantified from laminin-positive contours, and microvessel density from CD31-positive structures.

#### FST Gene Expression (RT-qPCR)

Total RNA was extracted from snap-frozen tissues using ExtractRNA reagent (Evrogen, Russia). In mice, injected and contralateral hindlimb muscles, liver, and gonads were sampled at necropsy. RNA quality was assessed by NanoDrop OneC, and Fst transgene expression was quantified by one-step SYBR Green RT-qPCR (OneTube RT-PCR mix, Evrogen) on a CFX96 system (Bio-Rad) using WPRE-specific primers with mouse β-actin as reference gene. Relative expression was calculated by the 2^(−ΔΔCt) method with the PBS group as calibrator. Primer sequences were as follows: mouse β-actin — Forward 5′-CATTGCTGACAGGATGCAGAAGG-3′, Reverse 5′-TGCTGGAAGGTGGACAGTGAGG-3′; FST transgene (WPRE-specific) — Forward 5′-TCCGGATCTTGCAACTCCAT-3′, Reverse 5′-CCACATAGCGTAAAAGGA GCA-3′.

#### FST Protein Quantification (ELISA)

Serum FST concentration and FST content in muscle lysates were measured using a commercial human Follistatin ELISA kit (SEKH-0183-96T, SolarBio, China) according to the manufacturer’s instructions. Muscle lysate values were normalised to total protein (BCA assay) and expressed as pg/mg total protein.

#### Statistical Analysis

For the primary endpoint of local muscle hypertrophy, we used a within-animal paired design (Festing, 2002): the right (treated) gastrocnemius was compared with the left (PBS, contralateral) gastrocnemius from the same animal by paired Student’s t-test, and the right/left ratio was calculated for each animal. Between-group comparison of right/left ratios used the Kruskal–Wallis test followed by post-hoc Mann–Whitney U with Bonferroni correction for three pairwise comparisons versus PBS (α = 0.0167). Cohen’s d was calculated as the within-animal effect size (Cohen, 2013; Lakens, 2013). Continuous safety endpoints (body weight, hematology, biochemistry) were compared between groups by one-way ANOVA with Tukey HSD or Kruskal–Wallis where the normality assumption was violated. Two-tailed p < 0.05 was considered significant. Data are presented as mean ± standard deviation unless otherwise indicated. Analyses were performed in Python (SciPy 1.13).

## Results

### General Health, Body Weight and Safety

All 36 mice completed the study without unscheduled deaths. Body weight increased progressively in all groups and did not differ between any treatment group and PBS at Days 32 or 39 (all p > 0.05). Haematology at Days 32, 55 and 115 showed no treatment-related changes in WBC, RBC, Hgb, Hct, or platelets. An isolated statistically significant decrease in mean platelet volume (MPV) was noted in AAV-FST males at Day 55 (6.1 ± 0.8 fL vs. PBS 8.2 ± 1.5 fL; p = 0.038), and an isolated increase at Day 115 (7.3 ± 1.0 fL vs. PBS 5.2 ± 0.8 fL; p = 0.016), without corroborating changes in platelet count, PCT, or PDW, and absent in females and the Combo group.

Liver enzymes, renal markers, and metabolic parameters were within normal ranges at all time points in both sexes (all p > 0.05; Figure 1). No statistically significant differences were detected among treatment groups for any parameter at any time point (Kruskal-Wallis test, p>0.05), with the exception of total bilirubin in males at day 32, which showed a significant group effect by Kruskal-Wallis (p=0.039); however, pairwise comparisons with the PBS control group did not reach significance (p>0.05 for all comparisons).

**Figure 1.**
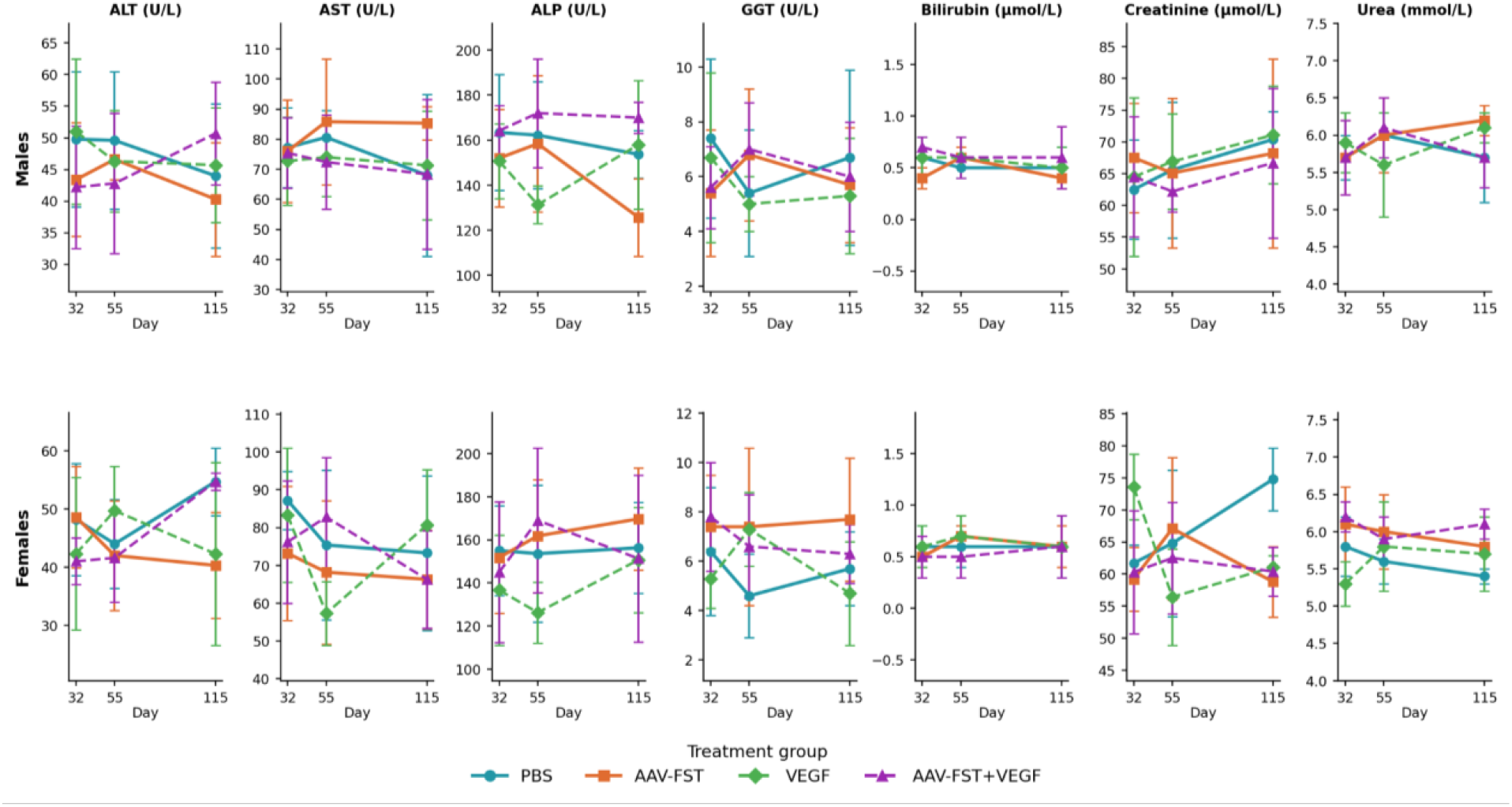
Serum biochemistry parameters in mice following administration of AAV-Follistatin, VEGF-plasmid, or their combination. Results are presented for males (upper panels) and females (lower panels) at days 32, 55, and 115 post-injection. Data are shown as mean ± SD. ALT, alanine aminotransferase; AST, aspartate aminotransferase; ALP, alkaline phosphatase; GGT, gamma-glutamyltransferase.

### FST Transgene Expression and Protein Levels

In mice at Day 55, AAV-FST monotherapy induced robust FST mRNA in the liver (~1,930-fold vs. PBS; p = 0.005) and injected muscle (~125-fold; p = 0.085). The Combo group showed comparable hepatic expression (~1,348-fold) but markedly higher muscle expression (~231-fold; p = 0.003), yielding a muscle-to-liver ratio 2.6-fold higher than monotherapy (0.171 vs. 0.065; Figure 2A). Gonadal FST expression was near background in the AAV-FST monotherapy group (~1.6-fold; p > 0.05) but was statistically significantly elevated in the Combo group (~4.7-fold; p = 0.032), without corroborating histopathological changes. At Day 115, FST mRNA remained markedly elevated in injected muscle in the AAV-FST group (~58-fold vs. PBS; p < 0.05) and was further increased in the Combo group (~122-fold vs. PBS; p < 0.05), whereas the VEGF plasmid alone showed no detectable increase in FST expression (dCt indistinguishable from PBS). Thus, at this later time point, the Combo maintained approximately 2.1-fold higher muscle FST mRNA than AAV-FST monotherapy, consistent with a sustained advantage in local transgene expression (Figure 2B).

**Figure 2.**
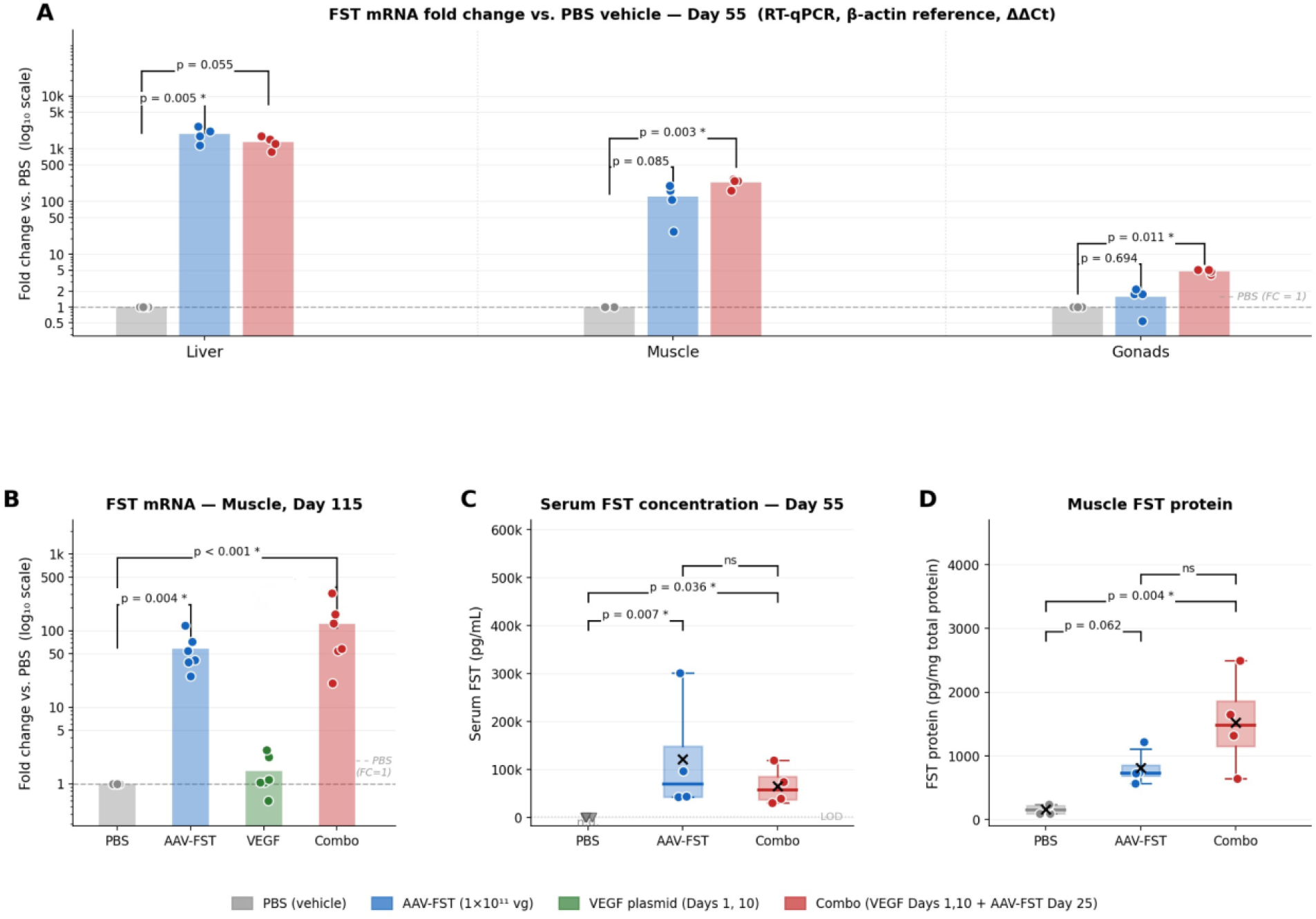
FST transgene expression assessed by RT-qPCR and ELISA in mice. (A) FST mRNA fold change vs. PBS in liver, skeletal muscle, and gonads at Day 55 by RT-qPCR (ΔΔCt method, β-actin reference gene). (B) FST mRNA fold change vs. PBS in injected hindlimb muscle at Day 115 by RT-qPCR (β-actin reference gene); VEGF plasmid group included as additional control. In panels A–B, bars represent group means, dots show individual animals (n = 4/group, Day 55; n = 6/group, Day 115), and the dashed line indicates PBS reference (fold change = 1.0). (C) Serum FST concentration in mice at Day 55 by ELISA (pg/mL, corrected for 1:200 dilution); PBS controls were below the assay detection limit (n.d., ▽). (D) FST protein in hindlimb muscle lysate normalised to total protein (pg/mg). In panels C–D, boxes represent median ± IQR, whiskers 1.5×IQR, × denotes the mean, and dots show individual animals (n = 4/group). ^*^ p < 0.05 vs. PBS; ns = not significant (Mann–Whitney test, Bonferroni correction).

At the protein level, serum FST was undetectable in PBS controls and elevated in both treated groups (p ≈ 0.021); the Combo group showed lower and less variable systemic levels (median 56,950 vs. 70,300 pg/mL; p = 0.49 vs. monotherapy). Muscle lysate FST was higher in the Combo group (median ~1,484 vs. ~733 pg/mg total protein), consistent with the mRNA data, though not statistically significant (p = 0.062 for AAV-FST vs. PBS; p = 0.004 for Combo vs. PBS; Figure 2C–D).

### Local muscle hypertrophy

Contralateral (PBS) limb mass was homogeneous across groups (Kruskal–Wallis H = 2.41, p = 0.492). In the sexes-pooled analysis (n = 6/group), the R/L ratio was 0.985 ± 0.048 (PBS), 1.113 ± 0.126 (AAV-FST), 0.980 ± 0.034 (VEGF alone), and 1.176 ± 0.091 (Combo) (Figure 3A). The Combo produced a highly significant within-animal effect (Δ = +0.240 g; paired t t(5) = 4.99, p = 0.004; Cohen’s d = 2.04); AAV-FST alone was borderline (p = 0.050; d = 1.05); VEGF alone did not differ from vehicle (p = 0.255). Between-group Kruskal–Wallis: H = 14.95, p = 0.0019; post-hoc Combo vs. PBS: p = 0.0022 (survives Bonferroni, α = 0.0167). Comparison of absolute right-limb mass (Figure 3B) showed a similar pattern (Kruskal–Wallis p = 0.006); pairwise p-values vs. PBS (AAV-FST p = 0.041; Combo p = 0.026) did not survive Bonferroni correction, making the R/L ratio the more sensitive primary endpoint. In males (n= 3/group), the Combo produced a statistically significant within-animal effect (R/L 1.213 ± 0.059; d = 3.24; p = 0.030); females showed consistent positive trends (d > 1.2) without reaching significance at n = 3. At the whole-cohort level (n = 6/group, 3 males and 3 females), the Combo group showed a uniform hypertrophic response, with all animals exhibiting R/L > 1.0 (6/6 “responders”) (Figure 3C). In contrast, AAV-FST monotherapy produced R/L > 1.0 in 5/6 animals, with one apparent non-responder (R/L = 0.97), whereas the VEGF plasmid group showed only 1/6 animals with R/L > 1.0, consistent with the absence of a meaningful effect.

**Figure 3.**
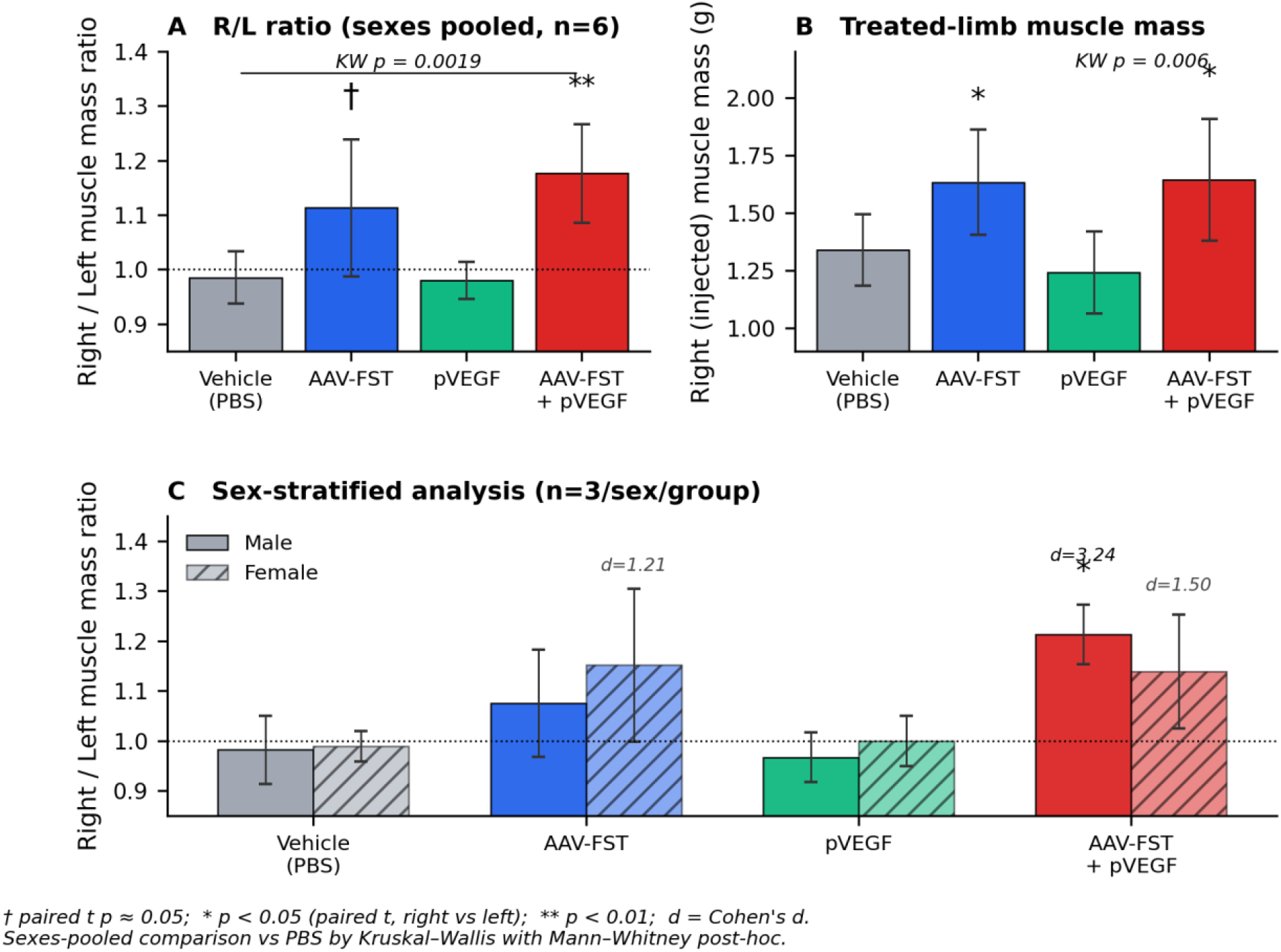
Local muscle hypertrophy after intramuscular AAV-FST and combined therapy. (A) Right (treated) /left (PBS contralateral) gastrocnemius mass ratio at Day 115 in mice (sexes pooled, n = 6 per group); dotted line indicates parity (R = L). Combination produced a significant within-animal effect (paired t-test p = 0.004); Kruskal–Wallis between-groups p = 0.0019. (B) Absolute right-limb (treated) muscle mass; Kruskal–Wallis p = 0.006; AAV-FST and combination both differ from PBS by Mann–Whitney post-hoc (p < 0.05). (C) Sex-stratified R/L ratios (n = 3 per sex per group). The largest and most reproducible effect was observed in the combination group in males (paired t-test p = 0.030; Cohen’s d = 3.24); both groups showed consistent positive trends in females (Cohen’s d > 1.2). † paired t p ≈ 0.05; ^*^ p < 0.05; ^**^ p < 0.01.

### Muscle Fibre Morphometry and Capillarisation

Laminin/CD31/DAPI co-staining demonstrated distinct and complementary effects of VEGF and follistatin on skeletal muscle architecture (Figure 4). VEGF-containing groups exhibited a visibly denser CD31^+^ microvascular network, whereas AAV-FST-treated muscles showed enlarged muscle fibres with preserved sarcolemmal integrity. The combination treatment produced both features simultaneously, with prominent fibre hypertrophy accompanied by increased local capillarisation.

**Figure 4.**
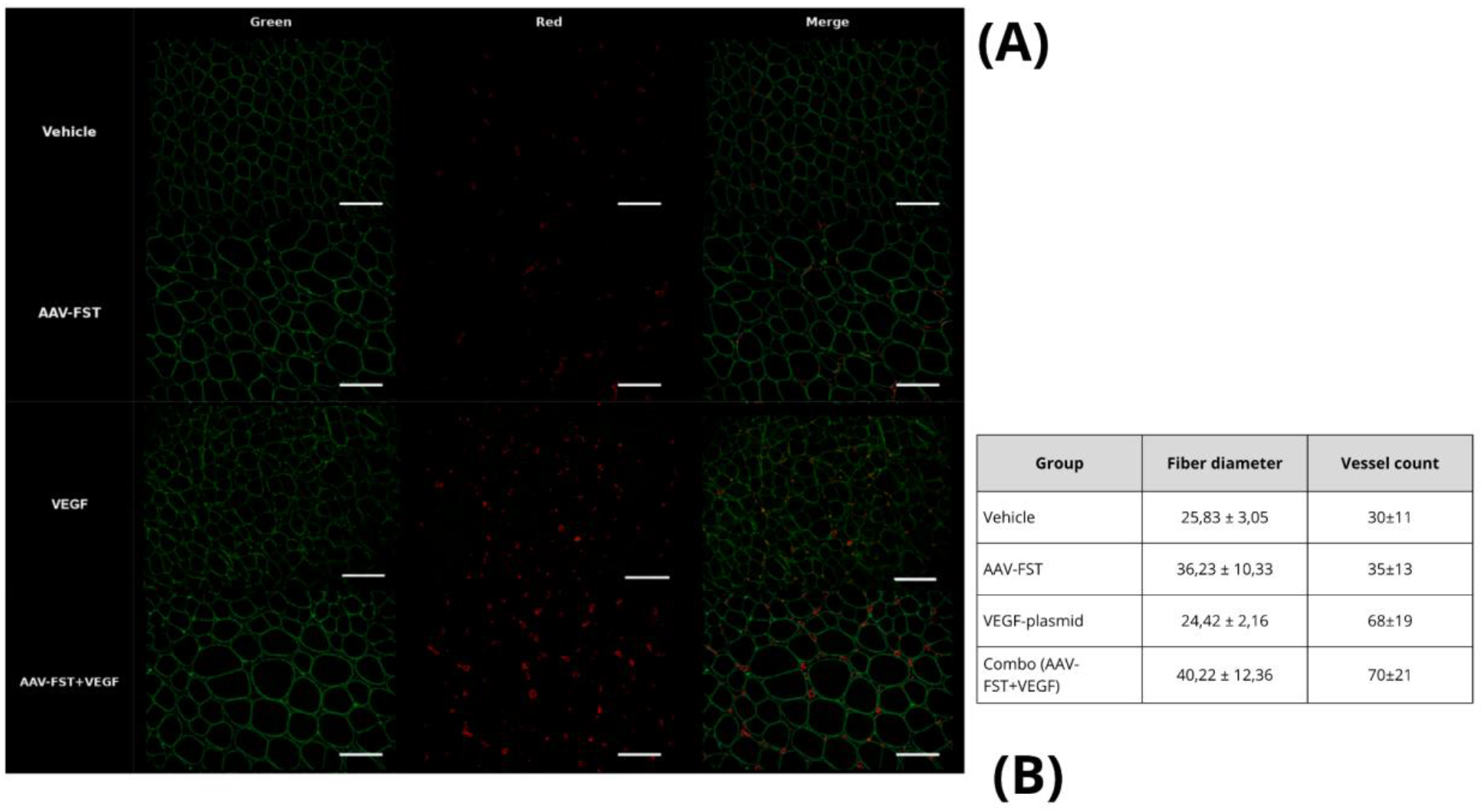
Laminin/CD31 immunohistochemistry and quantitative morphometry of injected hindlimb muscle at Day 115. **(A)** Representative transverse cryosections of the injected hindlimb muscle from Vehicle (PBS), AAV-FST, VEGF plasmid, and AAV-FST+VEGF (Combo) groups stained for laminin (green, sarcolemma) and CD31 (red, endothelial cells); nuclei are counterstained with DAPI (blue) in the merged channel. Columns correspond to single-channel laminin (Green), CD31 (Red), and merged images; rows correspond to treatment groups as indicated on the left. Scale bar = 50 µm. **(B)** Quantitative morphometry of muscle fibre diameter and CD31^+^ vessel counts per 20× field of view in the injected hindlimb muscle. Data are shown as mean ± SD.

Quantitative morphometry supported these observations. CD31^+^ vessel counts were approximately twofold higher in the VEGF plasmid (68 ± 19 vessels per 20× field) and Combo (70 ± 21) groups compared with Vehicle (30 ± 11) and AAV-FST monotherapy (35 ± 13), indicating a sustained angiogenic response associated with VEGF administration. In contrast, muscle fibre diameter was selectively increased in the AAV-FST (36.2 ± 10.3 μm) and Combo (40.2 ± 12.4 μm) groups relative to Vehicle (25.8 ± 3.1 μm) and VEGF monotherapy (24.4 ± 2.2 μm), with the largest fibres observed in the combination group.

### Histopathology and Organ Weights

Histopathological examination of the heart, lung, liver, kidney, brain, thymus, spleen, gonads, dorsal root ganglia, and injection-site skeletal muscle revealed no treatment-related pathological findings in any study group. Skeletal muscle architecture was preserved across all groups, with no evidence of necrosis, inflammatory infiltration, fibrosis, or central nucleation. Focal areas of reduced cytoplasmic eosinophilia were occasionally observed in vehicle-, AAV-FST-, and combination-treated muscles, whereas VEGF monotherapy showed more homogeneous staining (Supplementary Figures S1-S7). These findings were not associated with structural abnormalities and were considered non-adverse.

Organ weights measured at day 115 did not differ significantly between treatment groups in either sex (Kruskal–Wallis test, all p > 0.05; Supplementary Table S3). In particular, heart weight remained comparable across Vehicle, AAV-FST, VEGF, and combination-treated animals, with no evidence of cardiac enlargement or hypertrophy. Similarly, no treatment-related differences were observed in the weights of the liver, kidneys, lungs, spleen, thymus, brain, or gonads. As expected, increased mass was observed only in the treated hindlimb muscles of AAV-FST and combination-treated animals, consistent with the local hypertrophic effects of follistatin gene transfer.

## Discussion

The present study highlights three mechanistically linked findings. First, sequential intramuscular delivery of pl-VEGF165 followed by AAV9-FST344 produced greater and more reproducible local muscle hypertrophy than AAV-FST monotherapy, supporting the concept that angiogenic pre-conditioning can enhance the follistatin-mediated hypertrophic response. Second, this increase in muscle mass was accompanied by an approximately two-fold increase in CD31^+^ vessel counts, suggesting that the combination regimen promotes hypertrophy together with increased local capillarisation rather than hypertrophy alone. Third, VEGF pre-treatment was associated with a 2.6-fold higher muscle-to-liver transgene expression ratio, indicating a more favourable tissue expression profile following combination treatment. Although this difference did not reach formal statistical significance at the current sample size, it raises the possibility that VEGF-induced vascular remodelling may influence local AAV transduction efficiency and warrants further investigation. To our knowledge, this is the first study to combine AAV-delivered follistatin with a VEGF-A165 plasmid (Neovasculgen®), and the convergence of these findings supports the combination strategy as a biologically and translationally relevant platform for further development.

One possible explanation for the higher muscle-to-liver transgene expression ratio observed in the Combo group is VEGF-driven neovascularisation established during the 15-day interval between the final VEGF plasmid dose (Day 10) and AAV administration (Day 25). A denser local capillary network could increase the endothelial surface area accessible to the injected vector within the muscle interstitium, potentially enhancing local vector retention and increasing the probability of productive AAV–myofibre interactions. The kinetics of pl-VEGF165 expression are well suited to this sequential design: following intramuscular administration, VEGF mRNA peaks at Days 3–7 and becomes undetectable by approximately Day 35, while VEGF protein remains detectable for up to 35–50 days (Olea, 2009; Janavel, 2006). This expression window encompasses both VEGF plasmid administrations (Days 1 and 10) and the subsequent AAV injection (Day 25). Owing to its heparin-binding domain, VEGF165 signals through spatially restricted extracellular matrix-bound gradients that promote sprouting angiogenesis and pericyte recruitment, and has been shown to support the formation of functional microvascular networks in previous studies (Chen, 2010). Although the relationship between VEGF-induced capillarisation and subsequent AAV transduction efficiency was not directly tested in the present study, the observed increase in both CD31^+^ vessel density and muscle-targeted transgene expression suggests a potentially important interaction that warrants dedicated investigation.

A well-characterised limitation of myostatin/activin pathway blockade in isolation is capillary rarefaction. Soluble ActRIIB-Fc treatment increases muscle fibre size while reducing capillary density per fibre area and downregulating VEGF-A expression in skeletal muscle, whereas myostatin-null animals exhibit reduced specific force that has been attributed, at least in part, to inadequate capillarisation and impaired oxidative capacity (Hulmy, 2013; Matsakas, 2012). The VEGF co-delivery strategy employed in the present study was designed to address this potential limitation. Consistent with this rationale, VEGF monotherapy increased CD31^+^ vessel density without affecting muscle fibre size, whereas AAV-FST increased fibre diameter with minimal effect on capillarisation. Only the combination group exhibited both increased fibre size and increased vessel density, suggesting that the two interventions exert complementary and largely non-overlapping biological effects.

The biological relevance of this observation is supported by both experimental and clinical literature. Männistö et al. (2025) demonstrated that co-delivery of AAV-VEGF-B and a myostatin-propeptide vector rescued capillary rarefaction and enhanced muscle growth beyond myostatin inhibition alone, identifying endothelial cells and pericytes as key responders (Männistö, 2025). Similarly, Borselli et al. (2010) reported that combined VEGF and IGF-1 delivery promoted angiogenesis, reinnervation, and myogenesis more effectively than either factor alone in ischaemic skeletal muscle (Borselli, 2010). Human studies further support the relevance of capillarisation to muscle adaptation. Verdijk et al. (2016) observed that resistance training in older adults increased both type II fibre capillarisation and muscle fibre size, leading the authors to propose that improved capillary supply may facilitate the hypertrophic response in ageing muscle (Verdijk, 2016). Extending this observation, Snijders et al. (2017) found that baseline capillary-to-fibre ratio predicted the magnitude of hypertrophy following 24 weeks of resistance training (Snijders, 2017). Collectively, these findings suggest that vascular adaptation may represent an important determinant of the magnitude and quality of the hypertrophic response.

As muscle fibres enlarge, diffusion distances increase and metabolic demand rises, making vascular adaptation increasingly critical (Hellsten, 2025). Previous myostatin/activin-pathway interventions often increased muscle mass without improving physical performance, potentially owing to inadequate capillarisation. The simultaneous increase in fibre size and CD31^+^ vessel density in the Combo group may therefore represent a more physiologically integrated adaptation. Whether this translates into superior force generation or exercise capacity remains to be determined.

The combination was well tolerated, with no treatment-related clinical signs, histopathological abnormalities, or biochemical evidence of organ toxicity. A key advantage of using a plasmid for VEGF delivery is its transient, self-limiting expression profile, in contrast to AAV-VEGF, which can cause aberrant vascular growth; consistent with this, no vascular abnormalities were detected histologically (Karvinen, 2011; Gianni-Barrera, 2016). Previous clinical experience with pl-VEGF165 in peripheral arterial disease (Deev, 2015; Deev, 2017; Deev, 2018) further supports the translational feasibility of this approach.

Notably, despite detectable circulating FST protein and measurable transgene expression outside the injection site, hypertrophic effects remained predominantly localised to the treated hindlimb. No increase in heart weight, no histopathological abnormalities in cardiac tissue, and no evidence of generalized organ enlargement were observed. This dissociation between systemic exposure and local biological effect suggests that the current dosing regimen primarily drives regional muscle remodelling rather than widespread anabolic stimulation.

Several limitations should be acknowledged. The modest group size at the terminal timepoint (n = 3 per sex per group) limits statistical power for sex-stratified analyses. The study assessed local effects in the right hindlimb only; systemic bilateral delivery, which would be required for clinical sarcopenia or neuromuscular disease applications, was not evaluated. The follow-up period of 115 days does not address the long-term persistence of transgene expression relevant to adult gene therapy. Functional outcomes were not measured, and demonstrating that increased muscle mass translates into improved force production and functional performance will be an essential next step. In addition, CD31 staining provides evidence of increased vascular density but does not establish vessel maturity, perfusion, or functional oxygen delivery. Future studies should therefore incorporate direct assessments of muscle function, tissue perfusion, and oxidative capacity.

The proposed mechanism linking VEGF-induced capillarisation to enhanced local AAV transduction was not directly tested and remains inferential. Likewise, biodistribution assessment was limited to transgene expression analysis in a small number of tissues and did not include quantitative vector genome measurements. Although transgene expression was predominantly localised to the injected muscle, low-level gonadal expression was detected in the combination group, the biological significance of which remains uncertain. Future studies should incorporate comprehensive biodistribution analyses, including vector genome quantification across a broader range of tissues and expanded assessment of reproductive organs. Finally, the intervention should be validated in aged sarcopenic animals and disease-relevant neuromuscular models, where impaired vascular adaptation, fibrosis, and myostatin pathway dysregulation coexist.

## Conclusion

Sequential intramuscular delivery of pl-VEGF165 followed by AAV9-FST344 produced greater and more reproducible local muscle hypertrophy than AAV-FST monotherapy in C57BL/6 mice of both sexes, accompanied by increased CD31^+^ vessel density and a more favourable muscle-to-liver transgene expression profile. These findings provide proof-of-principle for a dual-vector strategy in which transient plasmid-mediated angiogenic stimulation enhances the local tissue response to durable AAV-delivered follistatin. Collectively, the results support further evaluation of this approach as a potential therapeutic platform for age-related muscle loss and neuromuscular disease.

## Acknowledgments

The authors thank the staff of Sechenov University for assistance with animal procedures, sample collection, and laboratory analyses conducted as part of this study. The authors also acknowledge the technical personnel involved in histological processing and molecular assays.

## Authors’ contributions

Anna Vakhrusheva conceived and designed the study, developed the scientific rationale, performed data analysis and interpretation, supervised the project, and wrote the original manuscript draft. Andrey A. Nedorubov conducted the animal experiments, sample collection, histological and molecular analyses, and contributed to data acquisition. Vladimir Leshko contributed to project administration, study coordination, and resource management. Ivan Morgunov contributed to funding acquisition, project administration, and resource provision. All authors critically reviewed the manuscript, contributed to its revision, and approved the final version for publication.

## Data Availability Statement

All data generated or analyzed during this study are included in this article and Supplementary Information. Further enquiries can be directed to the corresponding author.

## Supplementary

**Supplementary Table S1**. Body weight and body weight gain in C57BL/6 mice at Days 32 and 39 (means ± SD) and associated p values from Kruskal–Wallis tests.

**Supplementary Table S2**. Peripheral blood haematology in C57BL/6 mice after AAV-Follistatin and VEGF plasmid treatment.

**Supplementary Table S3**. Mean organ weights in C57BL/6 mice at Day 115, g.

**Supplementary Figure S1**. Histopathology of major organs in Vehicle-treated C57BL/6 mice at Day 55.

**Supplementary Figure S2**. Histopathology of major organs in AAV-Follistatin–treated C57BL/6 mice at Day 55.

**Supplementary Figure S3**. Histopathology of major organs in AAV-FST + VEGF–treated C57BL/6 mice at Day 55.

**Supplementary Figure S4**. Histopathology of major organs in Vehicle-treated C57BL/6 mice at Day 115.

**Supplementary Figure S5**. Histopathology of major organs in AAV-Follistatin–treated C57BL/6 mice at Day 115.

**Supplementary Figure S6**. Histopathology of major organs in VEGF-plasmid–treated C57BL/6 mice at Day 115.

**Supplementary Figure S7**. Histopathology of major organs in AAV-FST + VEGF–treated C57BL/6 mice at Day 115.

