## Supplementary for "Sequential VEGF-A165 plasmid and AAV-follistatin gene therapy enhances muscle hypertrophy and capillarisation in C57BL/6 mice"

**Supplementary Table S1.** Body weight and body weight gain in C57BL/6 mice at Days 32 and 39 (means ± SD) and associated p values from Kruskal–Wallis tests.

| **Sex** | **Measurement** | **Day** | **Vehicle (PBS)** | **AAV-FST (1×10¹¹ vg)** | **VEGF plasmid (100 µg)** | **AAV-FST + VEGF** | **p (Kruskal–Wallis)** |
| --- | --- | --- | --- | --- | --- | --- | --- |
| Male | Body weight, g | Day 0 | 20.60 ± 0.88 | 20.64 ± 0.98 | 20.38 ± 1.26 | 20.20 ± 0.70 | 0.828 |
|  |  | Day 32 | 32.66 ± 0.97 | 32.25 ± 1.92 | 32.64 ± 0.71 | 32.45 ± 1.32 | 0.993 |
|  |  | Day 39 | 35.81 ± 1.55 | 35.78 ± 1.69 | 35.99 ± 0.79 | 35.60 ± 1.28 | 0.979 |
|  | Body weight gain, % | Day 0 | 0 | 0 | 0 | 0 | – |
|  |  | Day 32 | 58.79 ± 8.84 | 56.55 ± 11.83 | 60.41 ± 6.96 | 60.82 ± 8.84 | 0.993 |
|  |  | Day 39 | 74.18 ± 11.98 | 73.60 ± 10.08 | 76.88 ± 7.41 | 76.44 ± 9.21 | 0.979 |
| Female | Body weight, g | Day 0 | 18.60 ± 0.50 | 18.81 ± 0.95 | 19.54 ± 0.39 | 19.27 ± 0.71 | 0.319 |
|  |  | Day 32 | 29.13 ± 0.36 | 29.31 ± 0.65 | 27.99 ± 0.74 | 28.46 ± 0.93 | 0.076 |
|  |  | Day 39 | 31.90 ± 0.83 | 31.54 ± 1.09 | 30.61 ± 1.03 | 31.08 ± 1.14 | 0.340 |
|  | Body weight gain, % | Day 0 | 0 | 0 | 0 | 0 | – |
|  |  | Day 32 | 41.63 ± 6.13 | 42.29 ± 7.67 | 37.54 ± 5.90 | 41.00 ± 6.64 | 0.076 |
|  |  | Day 39 | 55.07 ± 6.90 | 53.12 ± 9.52 | 50.36 ± 5.70 | 54.11 ± 9.69 | 0.340 |

Body weight gain calculated relative to Day 0 (baseline). No statistically significant differences between groups were detected at any time point (Kruskal–Wallis, all p > 0.05).

**Supplementary Table S2**. Peripheral blood haematology in C57BL/6 mice after AAV‑Follistatin and VEGF plasmid treatment.

| **32 day** | | | | | | | | |
| --- | --- | --- | --- | --- | --- | --- | --- | --- |
| **Male** | | | | | | | | |
|  | **Vehicle (PBS)** | | **AAV-Follistatin**  **(1*10^11 vg)** | | **VEGF plasmid (100 µg)**  **(100 мкг)** | | **AAV‑FST (day 25) + VEGF (days 1, 10)** | |
|  | **M** | **SD** | **M** | **SD** | **M** | **SD** | **M** | **SD** |
| **WBC** | 6,4 | 0,9 | 6,2 | 1,6 | 7,5 | 0,3 | 6,1 | 1,3 |
| **LYM%** | 82,9 | 1,3 | 82,2 | 2,5 | 82,4 | 2,4 | 81,4 | 1,2 |
| **MID%** | 12,0 | 1,3 | 12,7 | 1,4 | 12,9 | 1,5 | 12,2 | 1,5 |
| **GRAN%** | 5,1 | 1,1 | 5,1 | 1,7 | 4,6 | 1,3 | 6,4 | 0,7 |
| **LYM#** | 5,3 | 0,8 | 5,1 | 1,4 | 6,2 | 0,3 | 5,0 | 1,1 |
| **MID#** | 0,8 | 0,1 | 0,8 | 0,2 | 1,0 | 0,1 | 0,7 | 0,2 |
| **GRAN#** | 0,3 | 0,1 | 0,3 | 0,0 | 0,3 | 0,1 | 0,4 | 0,1 |
| **RBC** | 7,0 | 0,2 | 6,6 | 0,3 | 6,5 | 0,6 | 7,1 | 0,4 |
| **HGB** | 10,6 | 1,0 | 11,4 | 0,9 | 10,5 | 0,7 | 10,5 | 1,0 |
| **HCT** | 32,1 | 3,1 | 31,9 | 2,6 | 31,1 | 3,4 | 30,3 | 1,0 |
| **MCV** | 45,9 | 4,4 | 48,5 | 4,9 | 48,5 | 9,4 | 42,6 | 4,4 |
| **MCH** | 15,1 | 1,6 | 17,4 | 1,9 | 16,3 | 2,2 | 14,7 | 1,5 |
| **MCHC** | 33,1 | 3,7 | 36,1 | 4,9 | 34,0 | 4,0 | 34,7 | 3,6 |
| **RDW-SD** | 46,1 | 5,1 | 42,9 | 5,7 | 44,6 | 10,7 | 48,8 | 5,2 |
| **RDW-CV** | 21,5 | 2,0 | 20,4 | 2,0 | 20,7 | 4,2 | 23,2 | 2,1 |
| **PLT** | 352,4 | 52,7 | 332,2 | 46,4 | 300,7 | 37,5 | 350,0 | 39,6 |
| **MPV** | 8,0 | 1,1 | 6,2 | 1,5 | 7,2 | 2,2 | 6,5 | 1,7 |
| **PDW** | 6,8 | 0,4 | 6,7 | 0,3 | 6,7 | 0,1 | 6,4 | 0,2 |
| **PCT** | 0,3 | 0,0 | 0,2 | 0,1 | 0,2 | 0,1 | 0,2 | 0,0 |
| **P-LCR** | 9,4 | 1,0 | 10,0 | 1,6 | 10,4 | 2,2 | 8,6 | 2,1 |
| **Female** | | | | | | | | |
| **WBC** | 6,5 | 1,0 | 6,5 | 0,9 | 6,6 | 0,7 | 6,3 | 1,2 |
| **LYM%** | 83,8 | 1,2 | 83,7 | 1,8 | 81,4 | 0,6 | 81,9 | 1,7 |
| **MID%** | 11,6 | 0,7 | 11,7 | 1,2 | 13,0 | 1,7 | 13,0 | 0,8 |
| **GRAN%** | 4,6 | 1,0 | 4,6 | 1,5 | 5,6 | 1,3 | 5,1 | 1,6 |
| **LYM#** | 5,4 | 0,9 | 5,5 | 0,8 | 5,4 | 0,5 | 5,2 | 1,0 |
| **MID#** | 0,8 | 0,1 | 0,8 | 0,1 | 0,9 | 0,2 | 0,8 | 0,2 |
| **GRAN#** | 0,3 | 0,1 | 0,3 | 0,1 | 0,4 | 0,0 | 0,3 | 0,1 |
| **RBC** | 7,1 | 0,3 | 6,4 | 0,3 | 6,6 | 0,4 | 6,9 | 0,5 |
| **HGB** | 10,2 | 1,0 | 11,0 | 1,0 | 11,0 | 0,9 | 10,3 | 1,1 |
| **HCT** | 31,3 | 2,3 | 32,2 | 2,6 | 31,7 | 2,4 | 29,6 | 0,7 |
| **MCV** | 44,3 | 3,5 | 50,4 | 3,2 | 48,4 | 5,0 | 43,0 | 3,6 |
| **MCH** | 14,4 | 1,7 | 17,2 | 2,4 | 16,8 | 1,9 | 15,0 | 1,6 |
| **MCHC** | 32,7 | 4,8 | 34,3 | 5,4 | 34,7 | 0,3 | 34,9 | 4,4 |
| **RDW-SD** | 46,8 | 5,9 | 40,2 | 3,6 | 41,9 | 8,4 | 49,1 | 4,6 |
| **RDW-CV** | 22,2 | 1,6 | 19,5 | 1,2 | 20,4 | 2,2 | 22,9 | 1,9 |
| **PLT** | 326,0 | 69,5 | 355,0 | 91,5 | 330,0 | 70,9 | 364,0 | 71,7 |
| **MPV** | 7,3 | 1,2 | 6,9 | 2,7 | 6,6 | 1,4 | 7,0 | 2,4 |
| **PDW** | 6,4 | 0,4 | 7,0 | 0,3 | 6,3 | 0,1 | 6,3 | 0,3 |
| **PCT** | 0,2 | 0,0 | 0,2 | 0,0 | 0,2 | 0,0 | 0,2 | 0,1 |
| **P-LCR** | 8,7 | 2,0 | 6,8 | 2,1 | 8,6 | 4,2 | 10,1 | 3,5 |
| **55 day** | | | | | | | | |
| **Male** | | | | | | | | |
|  | **Vehicle (PBS)** | | **AAV-Follistatin**  **(1*10^11 vg)** | | **VEGF plasmid (100 µg)** | | **AAV‑FST (day 25) + VEGF (days 1, 10)** | |
|  | **M** | **SD** | **M** | **SD** | **M** | **SD** | **M** | **SD** |
| **WBC** | 6,2 | 1,3 | 5,9 | 1,5 | 6,7 | 1,7 | 5,8 | 1,2 |
| **LYM%** | 82,4 | 1,2 | 83,9 | 2,0 | 83,6 | 1,4 | 84,2 | 1,4 |
| **MID%** | 13,4 | 0,8 | 12,3 | 1,8 | 13,0 | 1,5 | 11,2 | 0,9 |
| **GRAN%** | 4,2 | 1,2 | 3,8 | 0,5 | 3,4 | 0,3 | 4,5 | 0,6 |
| **LYM#** | 5,1 | 1,1 | 4,9 | 1,2 | 5,6 | 1,4 | 4,9 | 1,0 |
| **MID#** | 0,8 | 0,2 | 0,7 | 0,3 | 0,9 | 0,3 | 0,7 | 0,1 |
| **GRAN#** | 0,3 | 0,1 | 0,2 | 0,0 | 0,2 | 0,0 | 0,3 | 0,1 |
| **RBC** | 7,0 | 0,3 | 6,7 | 0,5 | 6,6 | 0,3 | 6,8 | 0,6 |
| **HGB** | 9,7 | 0,6 | 10,9 | 0,7 | 10,8 | 0,8 | 10,5 | 1,0 |
| **HCT** | 31,2 | 2,4 | 30,7 | 1,4 | 32,5 | 1,1 | 29,9 | 2,2 |
| **MCV** | 44,5 | 3,1 | 45,9 | 3,8 | 49,4 | 2,3 | 44,6 | 5,4 |
| **MCH** | 13,9 | 0,9 | 16,3 | 1,8 | 16,4 | 0,7 | 15,6 | 1,8 |
| **MCHC** | 31,2 | 2,3 | 35,7 | 4,1 | 33,2 | 2,9 | 35,3 | 5,6 |
| **RDW-SD** | 46,3 | 0,5 | 44,6 | 2,8 | 39,3 | 3,1 | 46,6 | 5,4 |
| **RDW-CV** | 22,1 | 1,5 | 21,5 | 1,8 | 19,9 | 1,0 | 22,2 | 2,6 |
| **PLT** | 337,8 | 57,3 | 342,0 | 54,4 | 329,7 | 45,8 | 277,4 | 17,2 |
| **MPV** | 8,2 | 1,5 | 6,1 | 0,8 | 7,8 | 0,2 | 7,5 | 1,2 |
| **PDW** | 6,8 | 0,4 | 6,8 | 0,6 | 6,8 | 0,6 | 6,5 | 0,5 |
| **PCT** | 0,3 | 0,0 | 0,2 | 0,1 | 0,3 | 0,0 | 0,2 | 0,0 |
| **P-LCR** | 8,8 | 2,1 | 11,9 | 1,5 | 10,0 | 3,3 | 8,7 | 1,3 |
| **Female** | | | | | | | | |
| **WBC** | 6,6 | 1,2 | 6,4 | 1,1 | 5,9 | 0,7 | 6,3 | 0,9 |
| **LYM%** | 82,3 | 2,3 | 80,7 | 2,2 | 81,6 | 1,9 | 81,6 | 3,5 |
| **MID%** | 12,9 | 1,9 | 13,3 | 2,1 | 13,0 | 1,4 | 13,3 | 1,7 |
| **GRAN%** | 4,8 | 1,5 | 6,0 | 1,3 | 5,4 | 0,6 | 5,1 | 1,9 |
| **LYM#** | 5,4 | 1,1 | 5,2 | 1,0 | 4,8 | 0,6 | 5,1 | 0,7 |
| **MID#** | 0,8 | 0,1 | 0,8 | 0,1 | 0,8 | 0,1 | 0,8 | 0,2 |
| **GRAN#** | 0,3 | 0,1 | 0,4 | 0,1 | 0,3 | 0,0 | 0,3 | 0,1 |
| **RBC** | 6,9 | 0,4 | 6,6 | 0,5 | 6,4 | 0,3 | 6,8 | 0,5 |
| **HGB** | 10,3 | 1,1 | 10,6 | 0,9 | 10,0 | 1,6 | 10,1 | 0,8 |
| **HCT** | 31,9 | 2,1 | 31,5 | 1,5 | 32,0 | 2,0 | 31,9 | 2,7 |
| **MCV** | 46,2 | 4,3 | 47,9 | 4,2 | 50,2 | 5,2 | 47,2 | 1,2 |
| **MCH** | 14,9 | 2,1 | 16,0 | 1,3 | 15,7 | 2,1 | 15,0 | 1,1 |
| **MCHC** | 32,2 | 2,1 | 33,7 | 4,0 | 31,5 | 6,3 | 31,8 | 1,8 |
| **RDW-SD** | 42,7 | 3,4 | 43,3 | 4,5 | 41,3 | 2,4 | 45,4 | 2,2 |
| **RDW-CV** | 21,4 | 1,9 | 20,6 | 1,9 | 19,6 | 1,9 | 20,8 | 0,5 |
| **PLT** | 374,0 | 67,7 | 347,6 | 88,9 | 346,0 | 89,6 | 333,0 | 28,8 |
| **MPV** | 6,4 | 2,1 | 6,8 | 2,7 | 6,7 | 2,8 | 6,7 | 0,9 |
| **PDW** | 6,7 | 0,6 | 6,7 | 0,6 | 6,6 | 0,4 | 6,9 | 0,6 |
| **PCT** | 0,2 | 0,1 | 0,2 | 0,0 | 0,2 | 0,1 | 0,2 | 0,0 |
| **P-LCR** | 10,3 | 3,3 | 10,9 | 1,1 | 10,6 | 3,1 | 8,9 | 1,4 |
| **115 day** | | | | | | | | |
| **Male** | | | | | | | | |
|  | **Vehicle (PBS)** | | **AAV-Follistatin**  **(1*10^11 vg)** | | **VEGF plasmid (100 µg)** | | **AAV‑FST (day 25) + VEGF (days 1, 10)** | |
|  | **M** | **SD** | **M** | **SD** | **M** | **SD** | **M** | **SD** |
| **WBC** | 6,3 | 1,1 | 6,3 | 1,2 | 7,1 | 0,2 | 6,2 | 1,3 |
| **LYM%** | 82,0 | 2,1 | 82,1 | 0,4 | 82,4 | 1,5 | 82,6 | 3,0 |
| **MID%** | 12,0 | 1,4 | 12,0 | 0,6 | 12,1 | 1,4 | 13,0 | 1,7 |
| **GRAN%** | 6,0 | 1,5 | 5,9 | 0,6 | 5,5 | 1,4 | 4,4 | 1,7 |
| **LYM#** | 5,1 | 0,8 | 5,1 | 1,0 | 5,9 | 0,2 | 5,1 | 1,2 |
| **MID#** | 0,8 | 0,2 | 0,8 | 0,2 | 0,9 | 0,1 | 0,8 | 0,1 |
| **GRAN#** | 0,4 | 0,1 | 0,4 | 0,0 | 0,4 | 0,1 | 0,3 | 0,1 |
| **RBC** | 6,8 | 0,6 | 6,3 | 0,2 | 6,4 | 0,6 | 6,8 | 0,7 |
| **HGB** | 11,0 | 0,7 | 9,2 | 0,2 | 10,5 | 0,5 | 9,8 | 0,9 |
| **HCT** | 30,9 | 1,6 | 31,6 | 3,0 | 31,9 | 0,9 | 32,1 | 1,9 |
| **MCV** | 45,7 | 2,5 | 49,7 | 3,6 | 49,8 | 4,0 | 47,5 | 3,2 |
| **MCH** | 16,4 | 2,2 | 14,6 | 0,4 | 16,3 | 0,7 | 14,6 | 2,1 |
| **MCHC** | 35,8 | 4,0 | 29,4 | 2,6 | 32,8 | 1,3 | 30,7 | 2,5 |
| **RDW-SD** | 46,0 | 2,5 | 41,1 | 4,3 | 41,1 | 6,1 | 45,0 | 2,4 |
| **RDW-CV** | 21,5 | 1,2 | 19,8 | 1,5 | 19,8 | 1,6 | 20,7 | 1,4 |
| **PLT** | 422,3 | 13,6 | 355,3 | 57,0 | 405,7 | 47,2 | 371,3 | 97,6 |
| **MPV** | 5,2 | 0,8 | 7,3 | 1,0 | 6,4 | 0,3 | 6,4 | 0,2 |
| **PDW** | 6,6 | 0,6 | 6,7 | 0,5 | 7,4 | 0,1 | 7,2 | 0,2 |
| **PCT** | 0,2 | 0,0 | 0,3 | 0,0 | 0,3 | 0,0 | 0,2 | 0,1 |
| **P-LCR** | 11,1 | 0,8 | 9,1 | 3,7 | 9,9 | 1,6 | 9,7 | 2,7 |
| **Female** | | | | | | | | |
| **WBC** | 6,1 | 0,7 | 6,6 | 0,8 | 5,8 | 1,5 | 6,1 | 1,4 |
| **LYM%** | 81,5 | 1,5 | 83,4 | 2,0 | 83,0 | 1,0 | 84,7 | 1,3 |
| **MID%** | 13,5 | 1,1 | 11,3 | 1,7 | 12,2 | 1,8 | 11,0 | 0,3 |
| **GRAN%** | 5,0 | 0,7 | 5,4 | 1,2 | 4,8 | 1,5 | 4,2 | 1,2 |
| **LYM#** | 5,0 | 0,5 | 5,5 | 0,8 | 4,8 | 1,2 | 5,2 | 1,3 |
| **MID#** | 0,8 | 0,2 | 0,7 | 0,1 | 0,7 | 0,3 | 0,7 | 0,1 |
| **GRAN#** | 0,3 | 0,1 | 0,3 | 0,0 | 0,3 | 0,1 | 0,2 | 0,0 |
| **RBC** | 6,7 | 0,6 | 6,8 | 0,6 | 6,6 | 0,3 | 6,2 | 0,1 |
| **HGB** | 10,4 | 0,7 | 10,2 | 1,1 | 11,4 | 0,3 | 10,5 | 1,2 |
| **HCT** | 31,1 | 2,0 | 30,4 | 2,0 | 30,4 | 0,3 | 29,8 | 1,1 |
| **MCV** | 46,6 | 1,6 | 45,2 | 6,9 | 46,0 | 2,1 | 47,8 | 2,5 |
| **MCH** | 15,7 | 2,5 | 15,1 | 1,3 | 17,2 | 0,9 | 16,8 | 2,1 |
| **MCHC** | 33,6 | 4,4 | 33,8 | 4,5 | 37,4 | 1,4 | 35,1 | 3,7 |
| **RDW-SD** | 43,7 | 2,8 | 47,9 | 7,3 | 43,6 | 2,6 | 42,0 | 5,9 |
| **RDW-CV** | 21,1 | 0,7 | 22,0 | 3,4 | 21,3 | 1,0 | 20,5 | 1,1 |
| **PLT** | 298,3 | 1,2 | 399,0 | 14,1 | 325,3 | 43,4 | 349,0 | 89,4 |
| **MPV** | 9,3 | 0,9 | 5,6 | 1,4 | 7,0 | 1,6 | 5,8 | 3,0 |
| **PDW** | 6,8 | 0,6 | 6,8 | 0,2 | 7,0 | 0,8 | 6,4 | 0,4 |
| **PCT** | 0,3 | 0,0 | 0,2 | 0,0 | 0,2 | 0,0 | 0,2 | 0,0 |
| **P-LCR** | 6,1 | 0,4 | 8,4 | 1,4 | 8,1 | 2,5 | 9,7 | 2,9 |

Hematology parameters in mice at Days 32, 55, and 115 (means ± SD) and associated p values from Kruskal–Wallis tests and post‑hoc pairwise comparisons versus Vehicle (PBS). Data are shown for male and female C57BL/6 mice treated with Vehicle, AAV‑Follistatin (1×10¹¹ vg), VEGF plasmid (100 µg), or the combination regimen AAV‑FST (day 25) + VEGF plasmid (days 1 and 10). The table includes total and differential leukocyte counts (WBC, LYM, MID, GRAN; absolute and percentage values), erythrocyte indices (RBC, HGB, HCT, MCV, MCH, MCHC, RDW‑SD, RDW‑CV), and platelet parameters (PLT, MPV, PDW, PCT, P‑LCR). Global comparison of groups using the Kruskal–Wallis test revealed no statistically significant differences for any parameter in males at any time point (all p > 0.05). In females at Day 32, the Kruskal–Wallis test indicated significant group effects for RBC, MCV, RDW‑CV, and PDW; however, subsequent pairwise comparisons versus Vehicle did not identify any statistically significant differences (RBC: p = 0.055, 0.403, 1.000; MCV: p = 0.229, 1.000, 1.000; RDW‑CV: p = 0.229, 1.000, 1.000; PDW: p = 0.104, 1.000, 1.000 for AAV‑Follistatin, VEGF plasmid, and AAV‑FST+VEGF, respectively). Overall, no treatment‑related changes in peripheral haematology were detected.

**Supplementary Table S3.** Mean organ weights in C57BL/6 mice at Day 115, g.

| **Male** | | | | | | | | |
| --- | --- | --- | --- | --- | --- | --- | --- | --- |
|  | **Vehicle (PBS)** | | **AAV-Follistatin  (1*10^11 vg)** | | **VEGF plasmid (100 µg)** | | **AAV‑FST (day 25) + VEGF (days 1, 10)** | |
|  | **M** | **SD** | **M** | **SD** | **M** | **SD** | **M** | **SD** |
| **Heart** | 0,13 | 0,01 | 0,14 | 0,01 | 0,15 | 0,00 | 0,14 | 0,00 |
| **Thymus** | 0,05 | 0,01 | 0,05 | 0,00 | 0,05 | 0,01 | 0,04 | 0,01 |
| **Lung** | 0,28 | 0,02 | 0,28 | 0,02 | 0,30 | 0,03 | 0,260 | 0,01 |
| **Liver** | 2,45 | 0,13 | 2,43 | 0,12 | 2,44 | 0,14 | 2,39 | 0,15 |
| **Spleen** | 0,18 | 0,02 | 0,17 | 0,02 | 0,19 | 0,02 | 0,18 | 0,01 |
| **Kidney (2)** | 0,36 | 0,04 | 0,34 | 0,02 | 0,36 | 0,02 | 0,36 | 0,03 |
| **Left muscle** | 1,35 | 0,28 | 1,42 | 0,24 | 1,21 | 0,21 | 1,43 | 0,30 |
| **Right muscle** | 1,32 | 0,20 | 1,53 | 0,29 | 1,17 | 0,23 | 1,74 | 0,36 |
| **Brain** | 0,41 | 0,01 | 0,39 | 0,01 | 0,40 | 0,01 | 0,41 | 0,01 |
| **Female** | | | | | | | | |
|  | **Vehicle (PBS)** | | **AAV-Follistatin  (1*10^11 vg)** | | **VEGF-плазмида  (100 мкг)** | | **AAV‑FST (day 25) + VEGF (days 1, 10)** | |
|  | **M** | **SD** | **M** | **SD** | **M** | **SD** | **M** | **SD** |
| **Heart** | 0,13 | 0,01 | 0,14 | 0,02 | 0,13 | 0,00 | 0,14 | 0,01 |
| **Thymus** | 0,05 | 0,01 | 0,05 | 0,01 | 0,04 | 0,01 | 0,04 | 0,01 |
| **Lung** | 0,30 | 0,01 | 0,30 | 0,00 | 0,29 | 0,02 | 0,30 | 0,03 |
| **Liver** | 1,95 | 0,07 | 2,14 | 0,23 | 2,14 | 0,09 | 2,14 | 0,23 |
| **Spleen** | 0,17 | 0,01 | 0,19 | 0,01 | 0,16 | 0,03 | 0,17 | 0,01 |
| **Kidney (2)** | 0,31 | 0,03 | 0,30 | 0,04 | 0,32 | 0,04 | 0,26 | 0,01 |
| **Left muscle** | 1,38 | 0,11 | 1,53 | 0,28 | 1,31 | 0,07 | 1,38 | 0,22 |
| **Right muscle** | 1,36 | 0,14 | 1,73 | 0,11 | 1,31 | 0,12 | 1,55 | 0,13 |
| **Brain** | 0,40 | 0,01 | 0,41 | 0,01 | 0,39 | 0,01 | 0,40 | 0,01 |

Mean organ and hindlimb muscle weights (g) are shown for the Vehicle (PBS), AAV‑FST, VEGF plasmid, and AAV‑FST+VEGF groups (mean ± SD). Organs were weighed at terminal necropsy on Day 115. No significant between‑group differences in organ weights were detected by Kruskal–Wallis testing (p > 0.05 for all comparisons in both sexes).

**Supplementary Figure 1**. Histopathology of major organs in vehicle-treated C57BL/6 mice at Day 55.

| 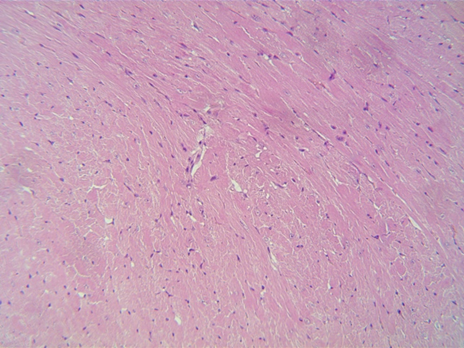 | 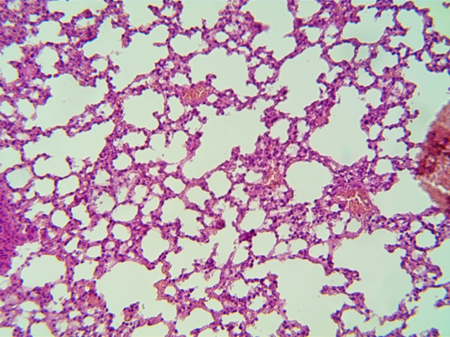 |
| --- | --- |
| Heart, H&E, ×200 (Vehicle, Day 55) | Lung, H&E, ×200 (Vehicle, Day 55) |
| 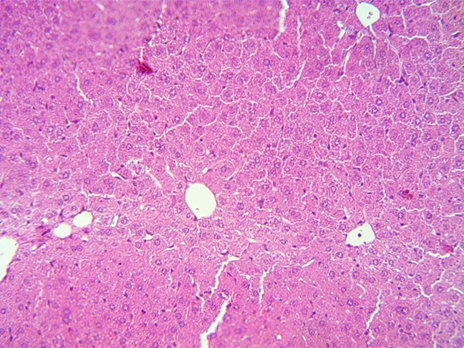 | 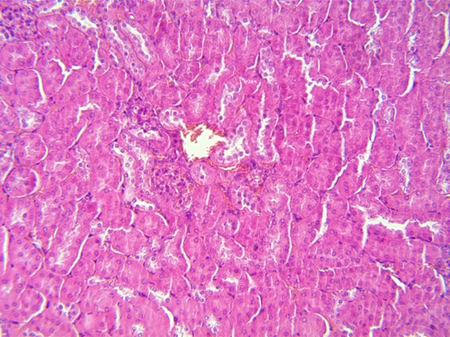 |
| Liver, H&E, ×200 (Vehicle, Day 55) | Kidney, H&E, ×200 (Vehicle, Day 55) |
| 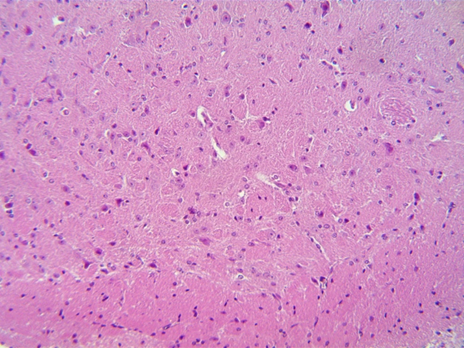 | 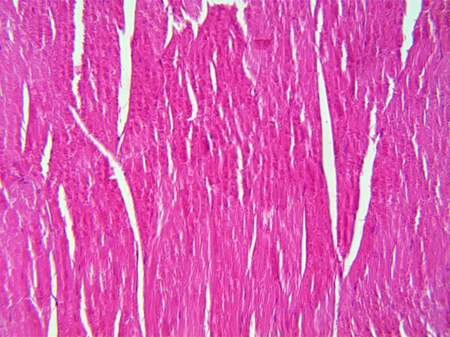 |
| Brain, H&E, ×200 (Vehicle, Day 55) | Skeletal muscle at injection site, H&E, ×200 (Vehicle, Day 55). |
| 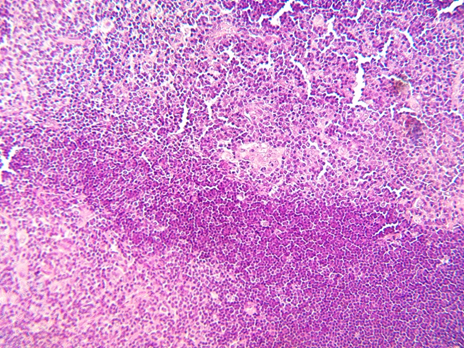 | 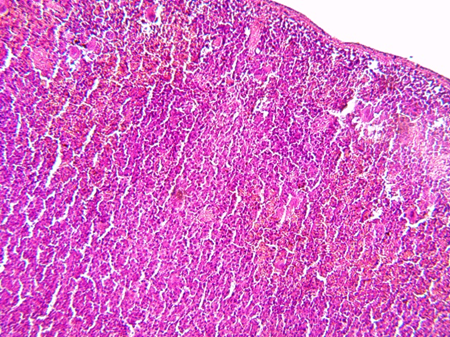 |
| Thymus, H&E, ×200 (Vehicle, Day 55) | Spleen, H&E, ×200 (Vehicle, Day 55) |
| 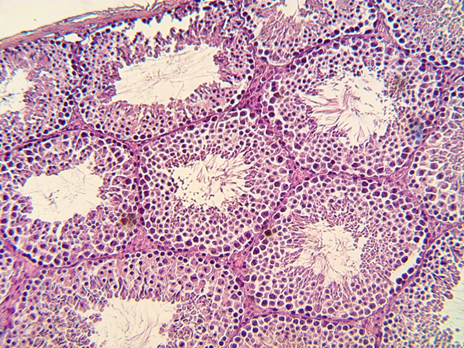 | 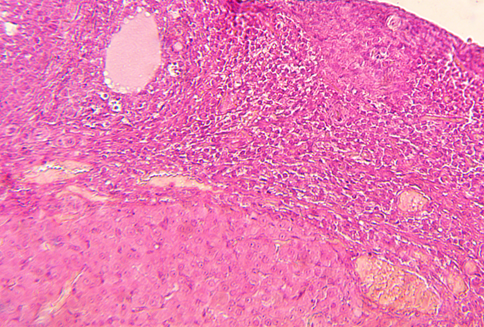 |
| Testis, H&E, ×200 (Vehicle, Day 55). | Ovary, H&E, ×200 (Vehicle, Day 55). |
| 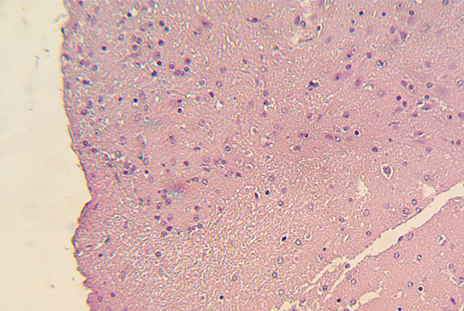 |  |
| Dorsal root ganglion, H&E, ×200 (Vehicle, Day 55) |  |

Representative H&E-stained sections (×200) from male and female C57BL/6 mice in the Vehicle (PBS) group show normal histological architecture of the heart, lung, liver, kidney, brain, injection-site skeletal muscle, thymus, spleen, testis, ovary with uterine tube, and dorsal root ganglion. The myocardium demonstrates preserved three-layer organisation with regular cardiomyocyte arrangement and intact interstitial vasculature; lung parenchyma shows well-aerated acini and bronchi with normal epithelial lining and only minimal perivascular and peribronchiolar lymphoid aggregates; liver displays a typical lobular pattern with regular hepatocyte plates, thin portal tracts and patent sinusoids; kidney presents normal cortico–medullary structure with intact glomeruli, tubules and collecting ducts; brain and cerebellum show preserved cortical layering and neuronal morphology; and skeletal muscle at the injection site consists of cross-striated fibres with peripheral nuclei and no inflammatory infiltrates or necrosis. Lymphoid organs (thymus, spleen) exhibit age-appropriate cortical–medullary differentiation and white/red pulp organisation, and gonads (testis, ovary) and dorsal root ganglia show normal cell composition and cytoarchitecture, with no treatment-related lesions detected in any examined tissue.

**Supplementary Figure 2**. Histopathology of major organs in AAV‑Follistatin–treated C57BL/6 mice at Day 55.

| 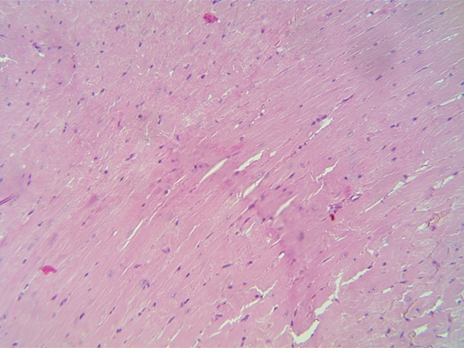 | 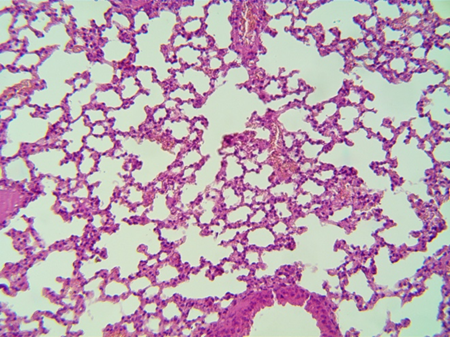 |
| --- | --- |
| Heart, H&E, ×200 (AAV‑Follistatin, Day 55) | Lung, H&E, ×200 (AAV‑Follistatin, Day 55) |
| 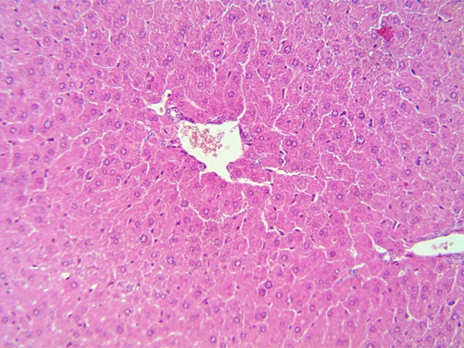 | 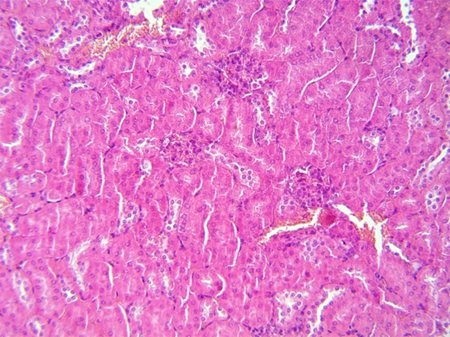 |
| Liver, H&E, ×200 (AAV‑Follistatin, Day 55) | Kidney, H&E, ×200 (AAV‑Follistatin, Day 55) |
| 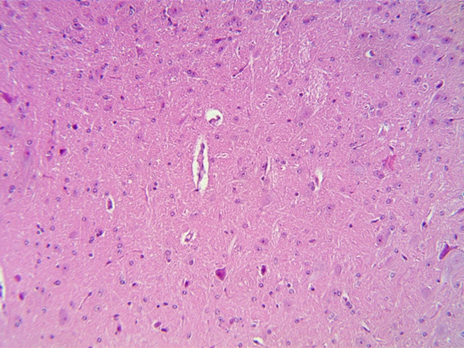 | 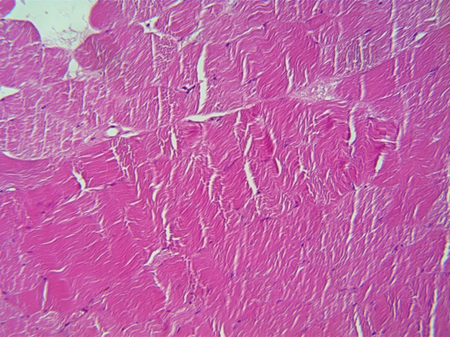 |
| Brain, H&E, ×200 (AAV‑Follistatin, Day 55) | Skeletal muscle at injection site, H&E, ×200 (AAV‑Follistatin, Day 55). |
| 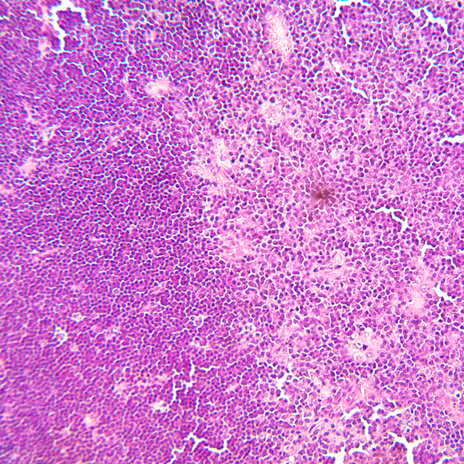 | 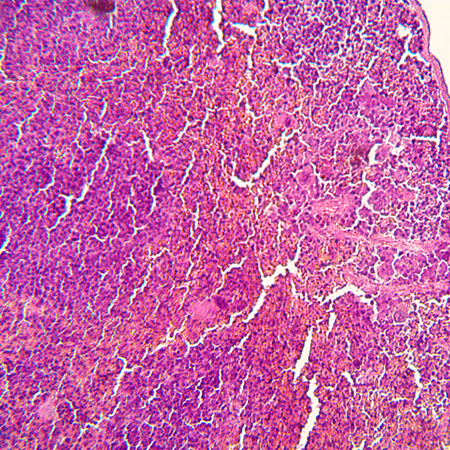 |
| Thymus, H&E, ×200 (AAV‑Follistatin, Day 55) | Spleen, H&E, ×200 (AAV‑Follistatin, Day 55) |
| 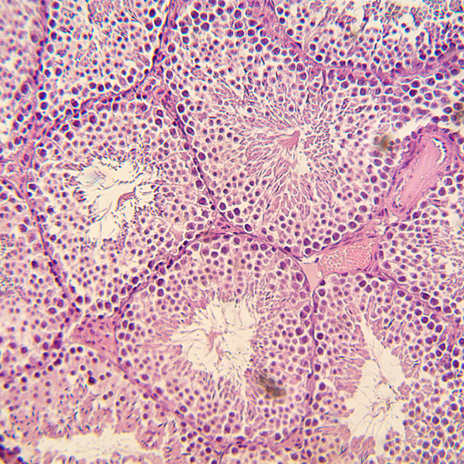 | 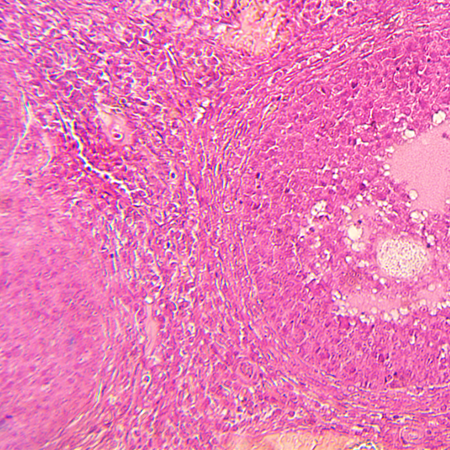 |
| Testis, H&E, ×200 (AAV‑Follistatin, Day 55). | Ovary, H&E, ×200 (AAV‑Follistatin, Day 55). |
| 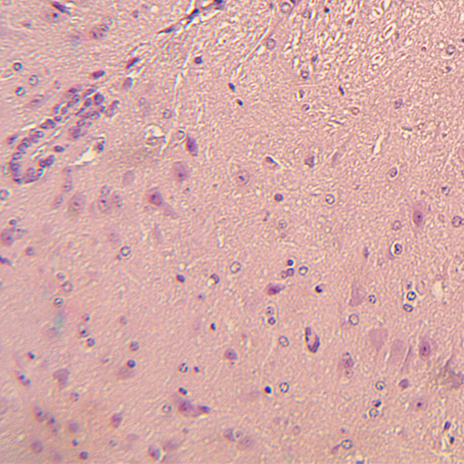 |  |
| Dorsal root ganglion, H&E, ×200 (AAV‑Follistatin, Day 55) |  |

Representative H&E‑stained sections (×200) from the AAV‑Follistatin (1×10¹¹ vg) group show normal histological architecture of the heart, lung, liver, kidney, brain, injection‑site skeletal muscle, thymus, spleen, gonads and dorsal root ganglia, with no treatment‑related lesions detected in any examined tissue.

**Supplementary Figure 3**. Histopathology of major organs in AAV‑FST + VEGF–treated C57BL/6 mice at Day 55.

| 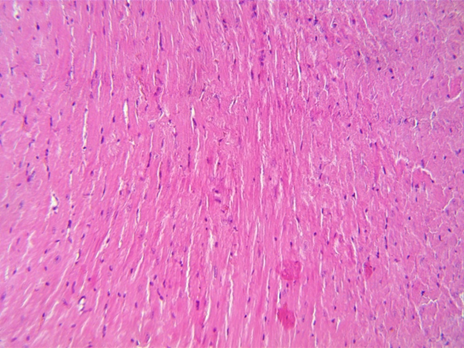 | 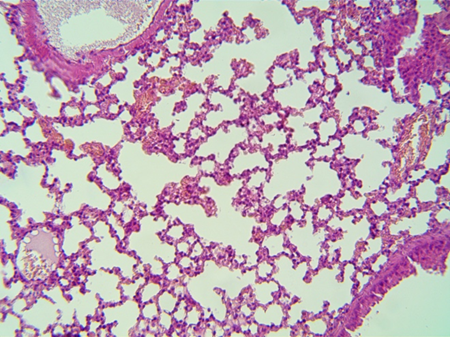 |
| --- | --- |
| Heart, H&E, ×200 (Combo, Day 55) | Lung, H&E, ×200 (Combo, Day 55) |
| 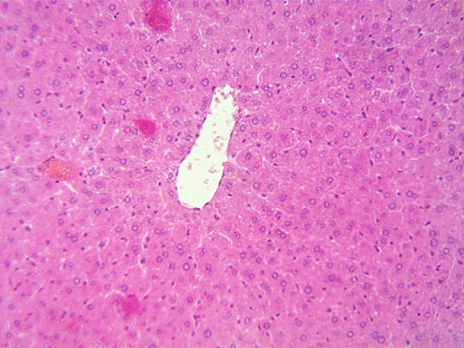 | 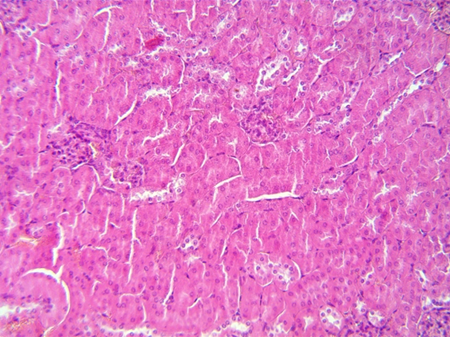 |
| Liver, H&E, ×200 (Combo, Day 55) | Kidney, H&E, ×200 (Combo, Day 55) |
| 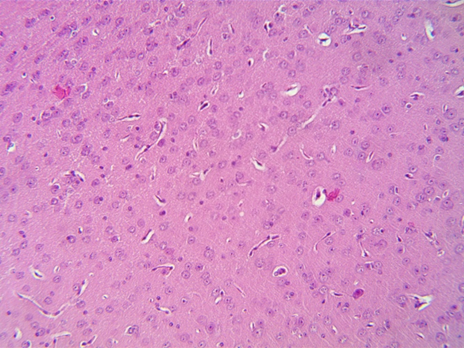 | 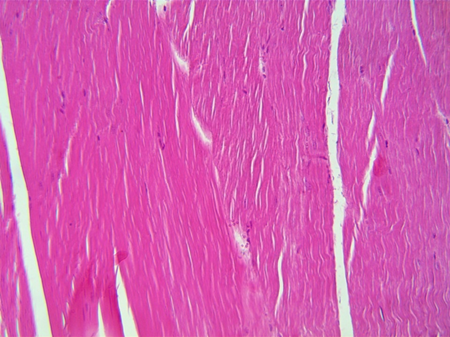 |
| Brain, H&E, ×200 (Combo, Day 55) | Skeletal muscle at injection site, H&E, ×200 (Combo, Day 55). |
| 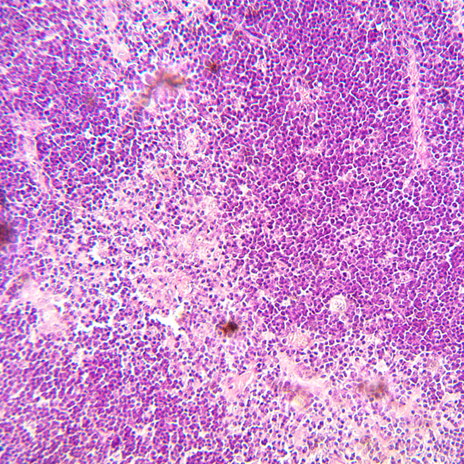 | 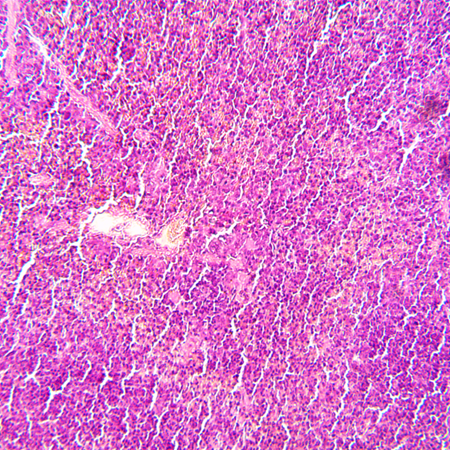 |
| Thymus, H&E, ×200 (Combo, Day 55) | Spleen, H&E, ×200 (Combo, Day 55) |
| 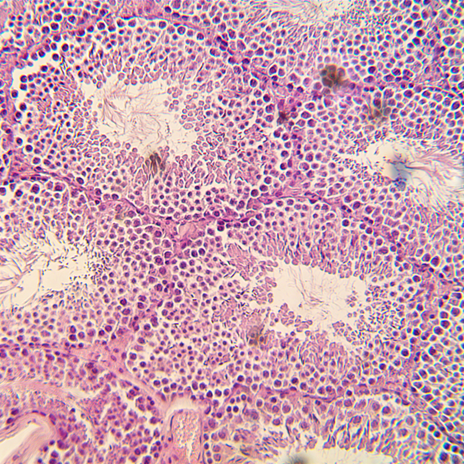 | 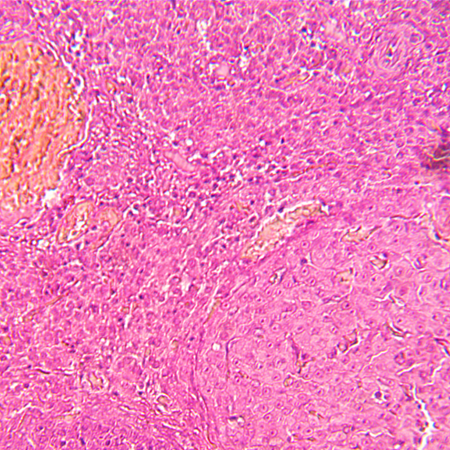 |
| Testis, H&E, ×200 (Combo, Day 55). | Ovary, H&E, ×200 (Combo, Day 55). |
| 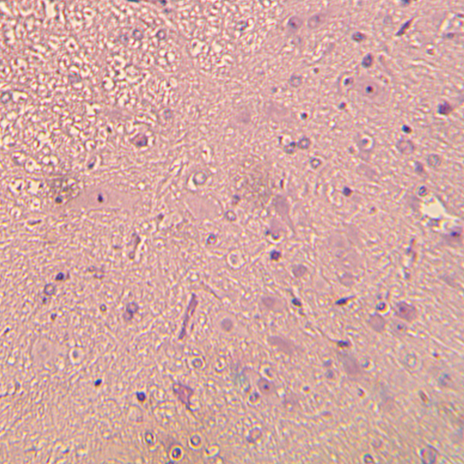 |  |
| Dorsal root ganglion, H&E, ×200 (Combo, Day 55) |  |

Representative H&E‑stained sections (×200) from the Combo (AAV‑FST (Day 25) + VEGF (Days 1 and 10)) group show normal histological architecture of the heart, lung, liver, kidney, brain, injection‑site skeletal muscle, thymus, spleen, gonads and dorsal root ganglia, with no treatment‑related lesions observed in any examined tissue.

**Supplementary Figure 4**. Histopathology of major organs in Vehicle‑treated C57BL/6 mice at Day 115.

| 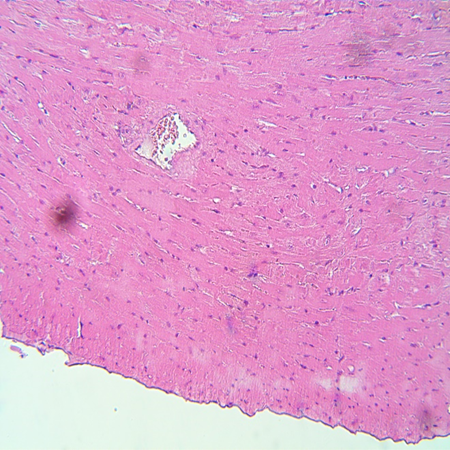 | 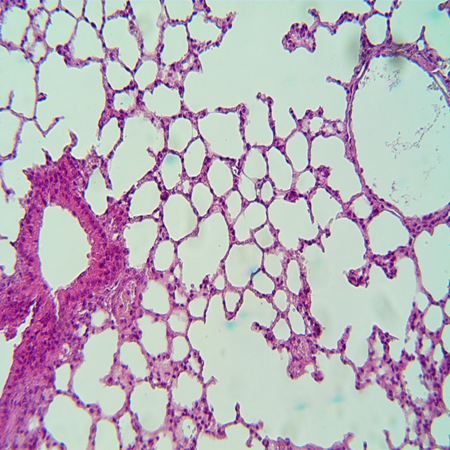 |
| --- | --- |
| Heart, H&E, ×200 (Vehicle, Day 115) | Lung, H&E, ×200 (Vehicle, Day 115) |
| 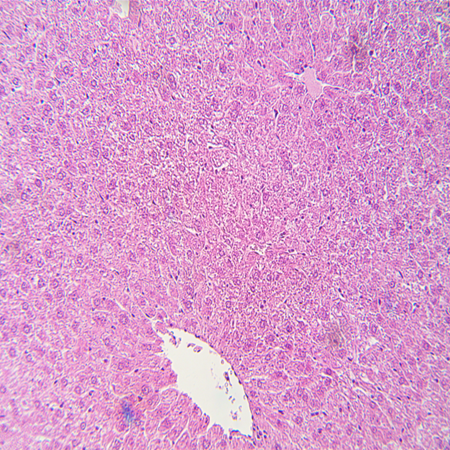 | 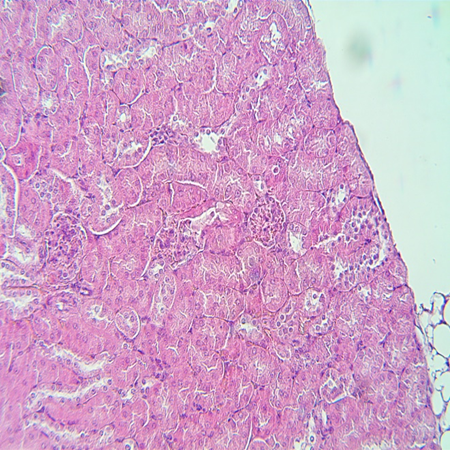 |
| Liver, H&E, ×200 (Vehicle, Day 115) | Kidney, H&E, ×200 (Vehicle, Day 115) |
| 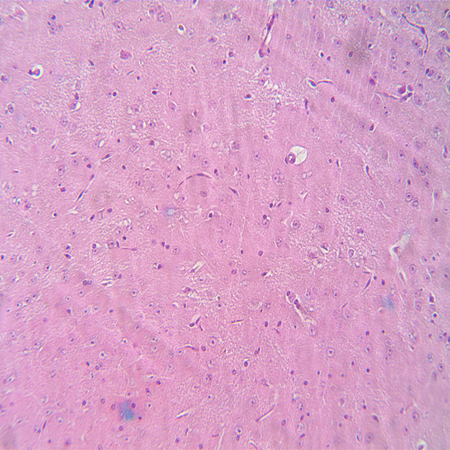 | 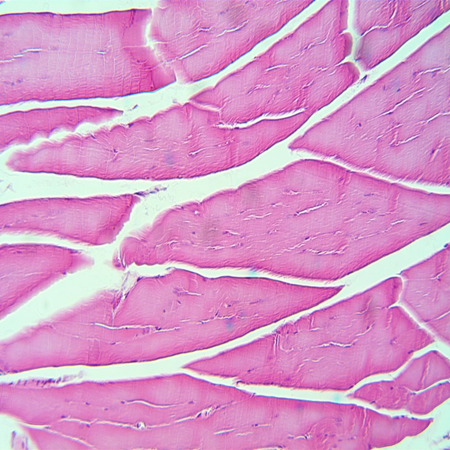 |
| Brain, H&E, ×200 (Vehicle, Day 115) | Skeletal muscle at injection site, H&E, ×200 (Vehicle, Day 115). |
| 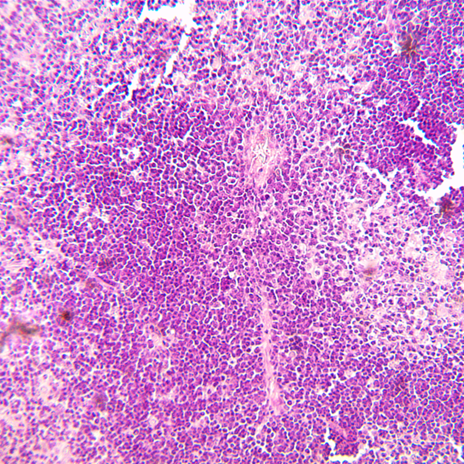 | 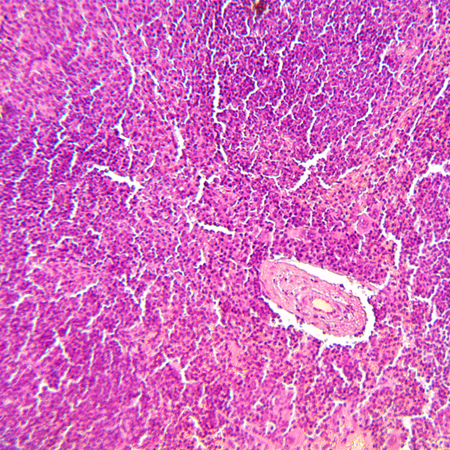 |
| Thymus, H&E, ×200 (Vehicle, Day 115) | Spleen, H&E, ×200 (Vehicle, Day 115) |
| 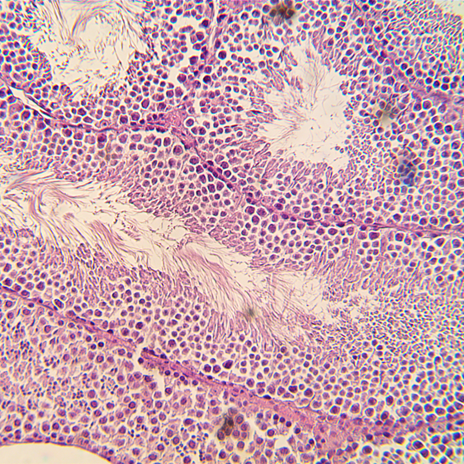 | 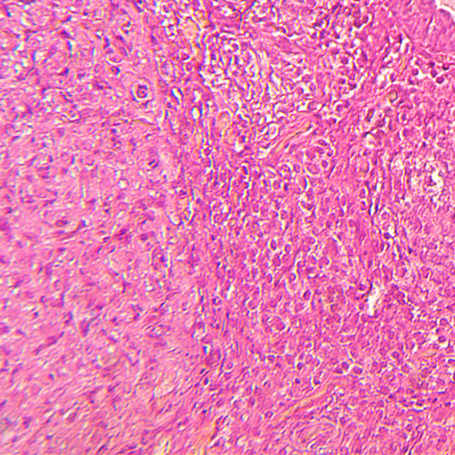 |
| Testis, H&E, ×200 (Vehicle, Day 115). | Ovary, H&E, ×200 (Vehicle, Day 115). |
| 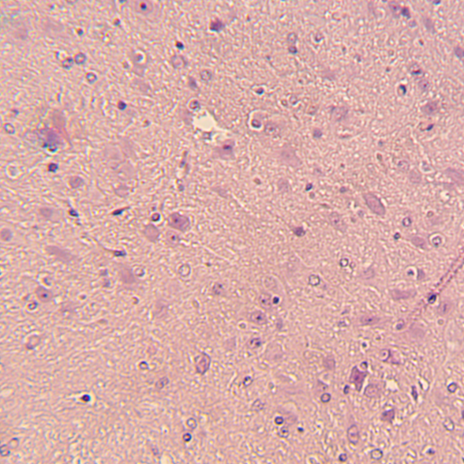 |  |
| Dorsal root ganglion, H&E, ×200 (Vehicle, Day 115) |  |

**Supplementary Figure 5**. Histopathology of major organs in AAV‑Follistatin–treated C57BL/6 mice at Day 115.

| 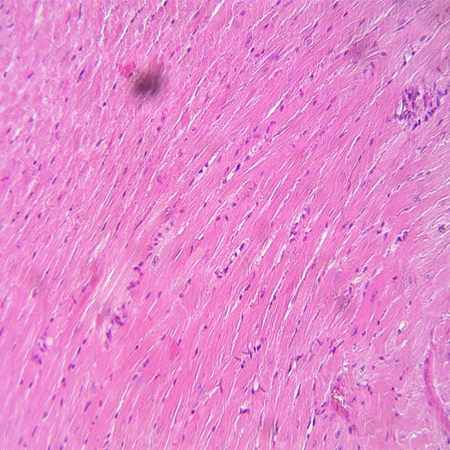 | 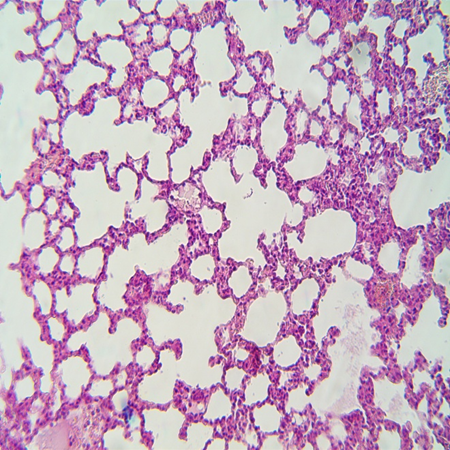 |
| --- | --- |
| Heart, H&E, ×200 (AAV‑Follistatin, Day 115) | Lung, H&E, ×200 (AAV‑Follistatin, Day 115) |
| 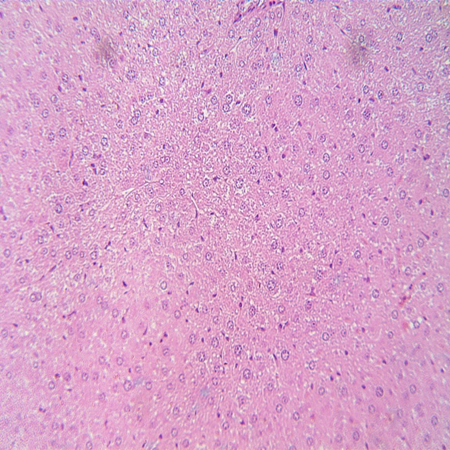 | 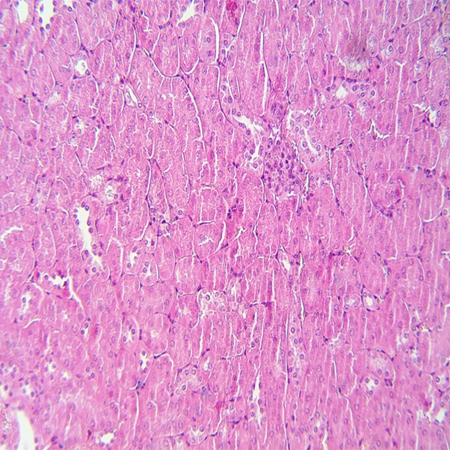 |
| Liver, H&E, ×200 (AAV‑Follistatin, Day 115) | Kidney, H&E, ×200 (AAV‑Follistatin, Day 115) |
| 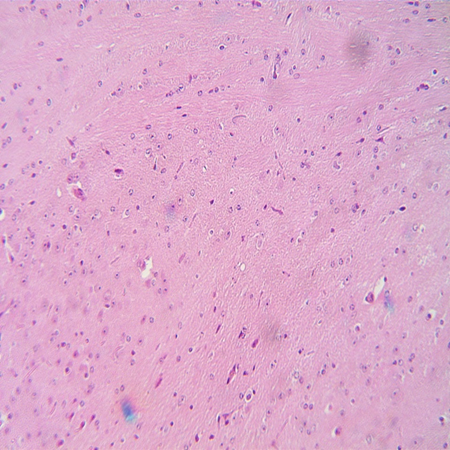 | 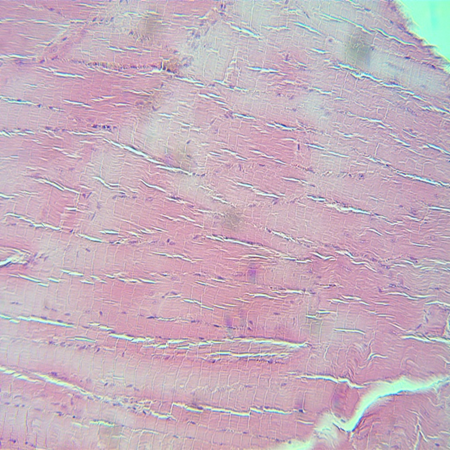 |
| Brain, H&E, ×200 (AAV‑Follistatin, Day 115) | Skeletal muscle at injection site, H&E, ×200 (AAV‑Follistatin, Day 115). |
| 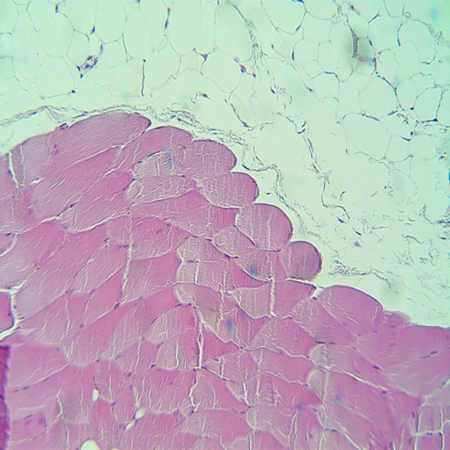 | 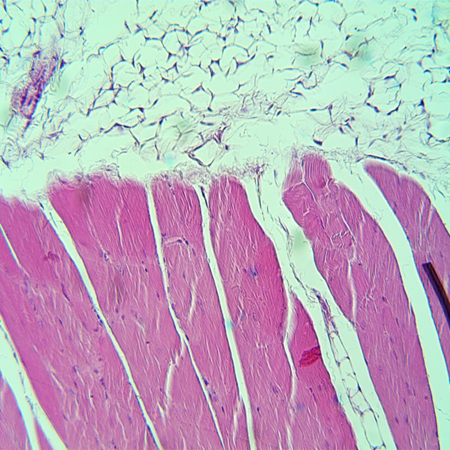 |
| Skeletal muscle at injection site, H&E, ×200 (AAV‑Follistatin, Day 115). | Skeletal muscle at injection site, H&E, ×200 (AAV‑Follistatin, Day 115). |
| 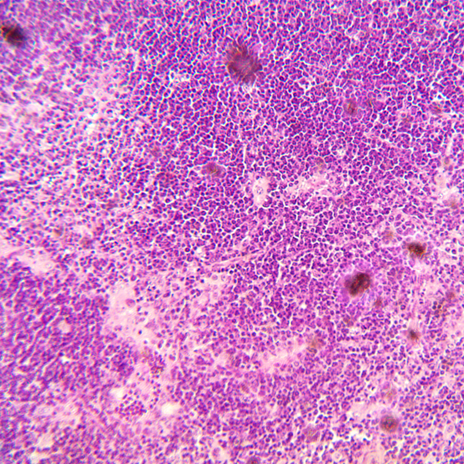 | 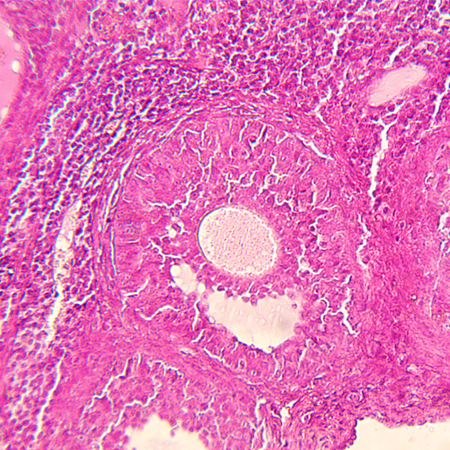 |
| Thymus, H&E, ×200 (AAV‑Follistatin, Day 115) | Spleen, H&E, ×200 (AAV‑Follistatin, Day 115) |
| 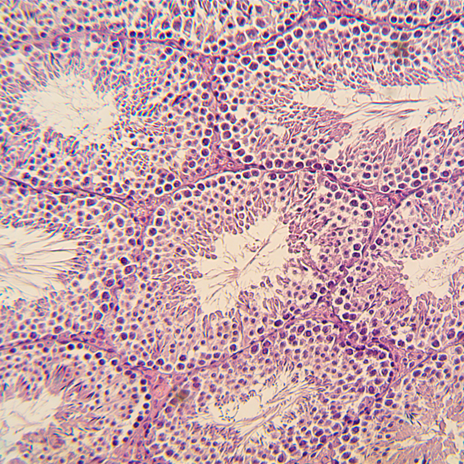 | 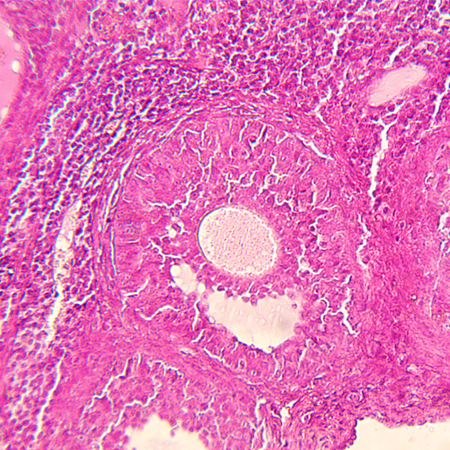 |
| Testis, H&E, ×200 (AAV‑Follistatin, Day 115). | Ovary, H&E, ×200 (AAV‑Follistatin, Day 115). |
| 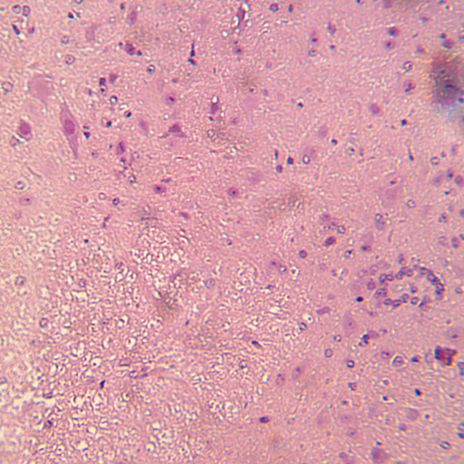 |  |
| Dorsal root ganglion, H&E, ×200 (AAV‑Follistatin, Day 115) |  |

Representative H&E‑stained sections (×200) from the AAV‑Follistatin (1×10¹¹ vg) group show normal histological architecture of the heart, lung, liver, kidney, brain, thymus, spleen, gonads and dorsal root ganglia; skeletal muscle at the injection site displays overall preserved morphology of cross‑striated fibres with peripheral nuclei, with focal areas of cytoplasmic clearing and prominent peri‑ and intramuscular adipose tissue, but no features of inflammatory or degenerative pathology.

**Supplementary Figure 6**. Histopathology of major organs in VEGF-plasmid–treated C57BL/6 mice at Day 115.

| 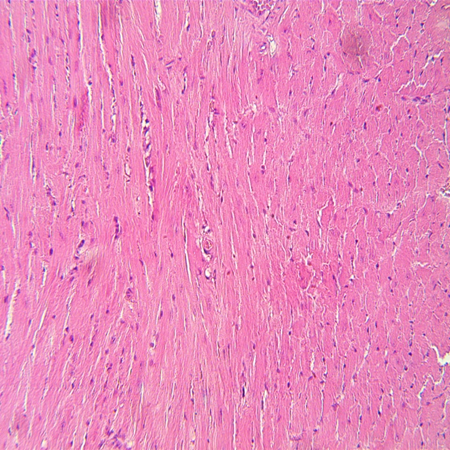 | 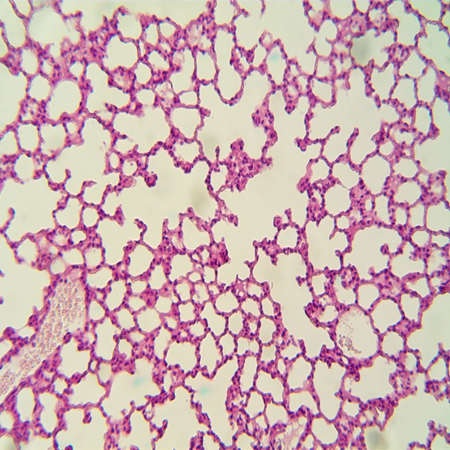 |
| --- | --- |
| Heart, H&E, ×200 (VEGF-plasmid, Day 115) | Lung, H&E, ×200 (VEGF-plasmid, Day 115) |
| 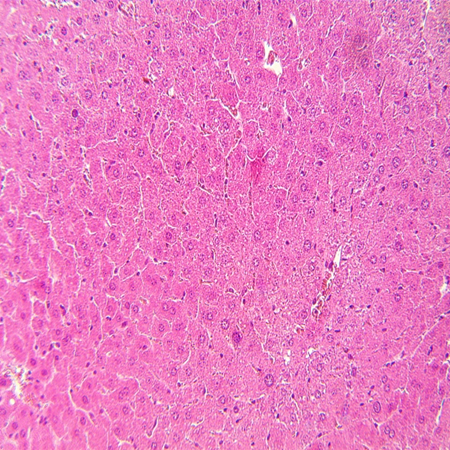 | 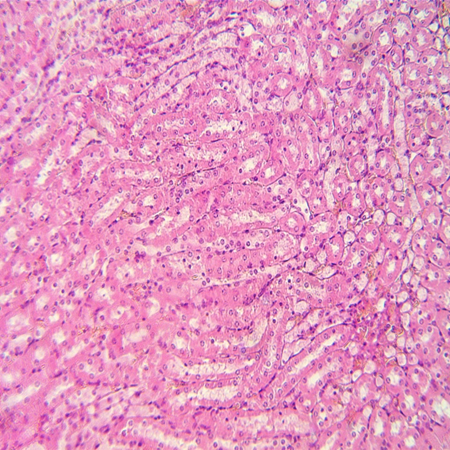 |
| Liver, H&E, ×200 (VEGF-plasmid, Day 115) | Kidney, H&E, ×200 (VEGF-plasmid, Day 115) |
| 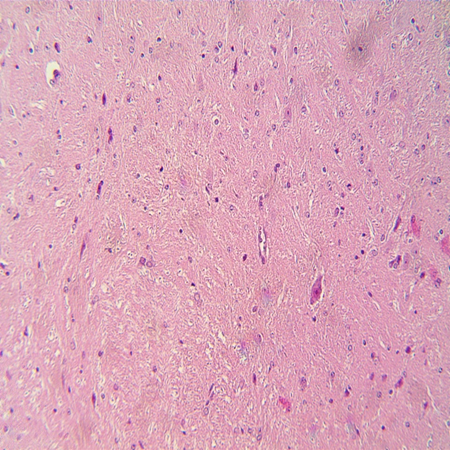 | 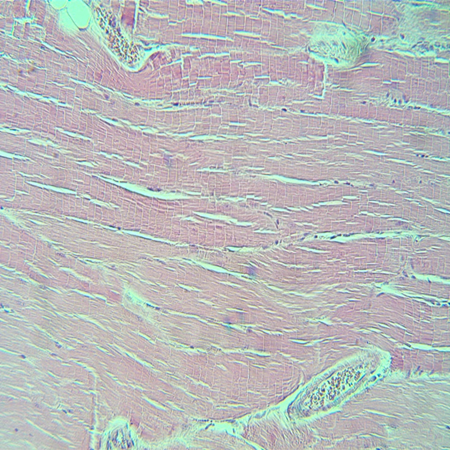 |
| Brain, H&E, ×200 (VEGF-plasmid, Day 115) | Skeletal muscle at injection site, H&E, ×200 (VEGF-plasmid, Day 115). |
| 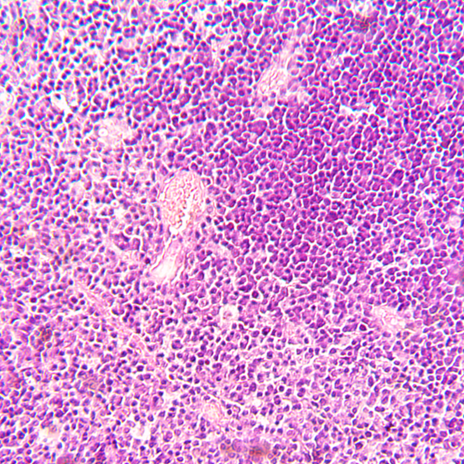 | 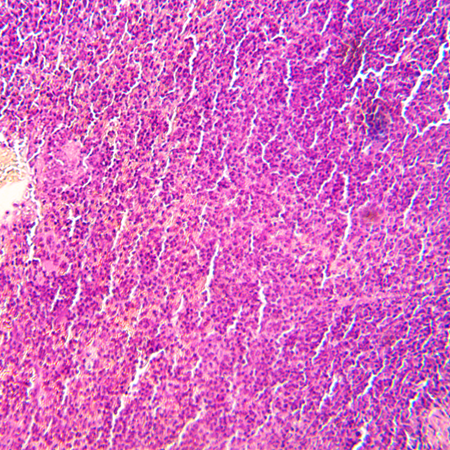 |
| Thymus, H&E, ×200 (VEGF-plasmid, Day 115) | Spleen, H&E, ×200 (VEGF-plasmid, Day 115) |
| 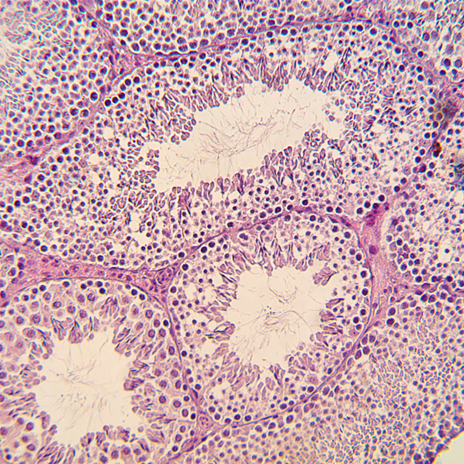 | 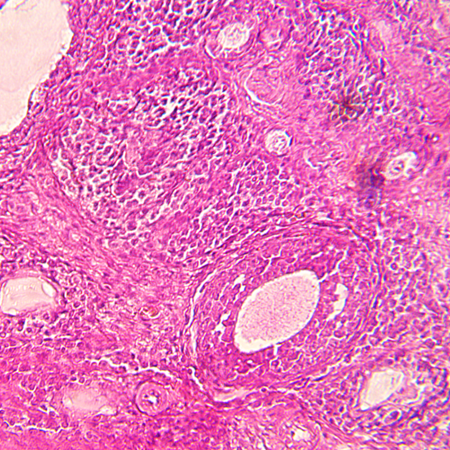 |
| Testis, H&E, ×200 (VEGF-plasmid, Day 115). | Ovary, H&E, ×200 (VEGF-plasmid, Day 115). |
| 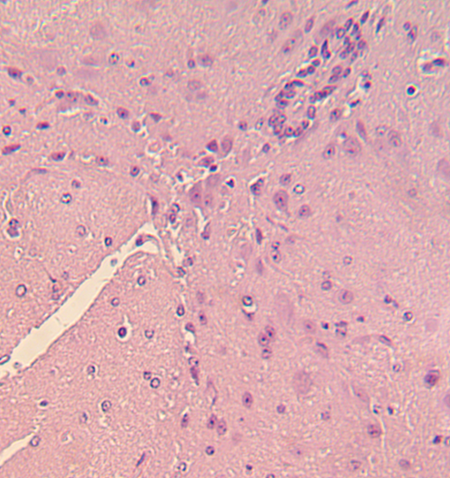 |  |
| Dorsal root ganglion, H&E, ×200 (VEGF-plasmid, Day 115) |  |

Representative H&E‑stained sections (×200) from the VEGF plasmid (100 µg) group show normal histological architecture of the heart, lung, liver, kidney, brain, thymus, spleen, gonads and dorsal root ganglia. Skeletal muscle at the injection site exhibits typical morphology of cross‑striated fibres with peripheral nuclei and uniformly stained cytoplasm, while intramuscular and perimysial blood vessels are readily visualised and appear more prominent than in Vehicle‑treated animals, without evidence of inflammatory or degenerative pathology.

**Supplementary Figure 7**. Histopathology of major organs in AAV‑FST + VEGF–treated C57BL/6 mice at Day 115.

| 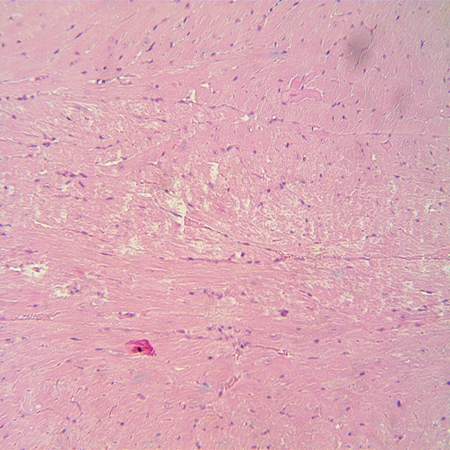 | 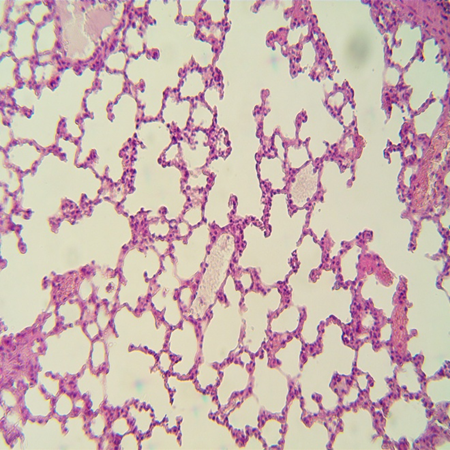 |
| --- | --- |
| Heart, H&E, ×200 (AAV‑FST + VEGF Day 115) | Lung, H&E, ×200 (AAV‑FST + VEGF Day 115) |
| 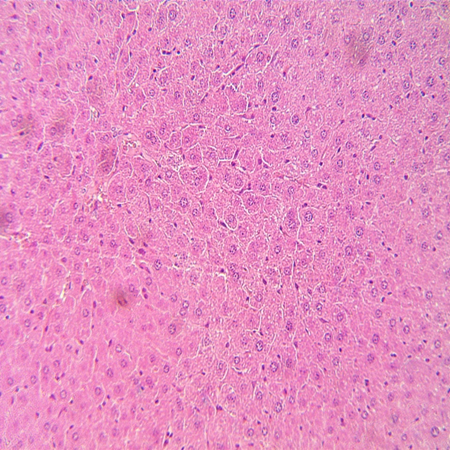 | 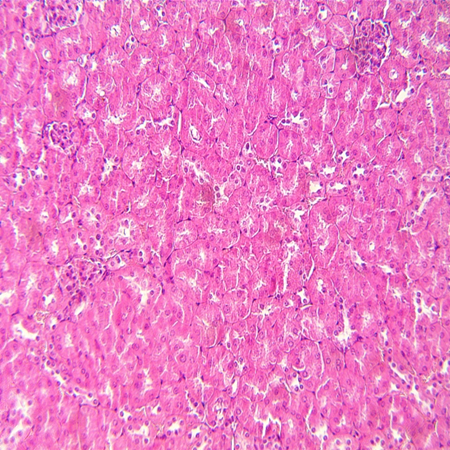 |
| Liver, H&E, ×200 (AAV‑FST + VEGF Day 115) | Kidney, H&E, ×200 (AAV‑FST + VEGF Day 115) |
| 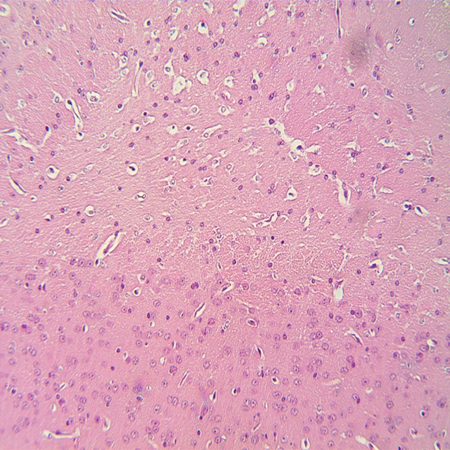 | 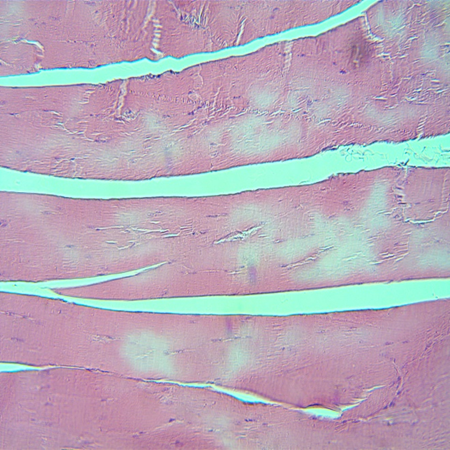 |
| Brain, H&E, ×200 (AAV‑FST + VEGF Day 115) | Skeletal muscle at injection site, H&E, ×200 (AAV‑FST + VEGF Day 115). |
| 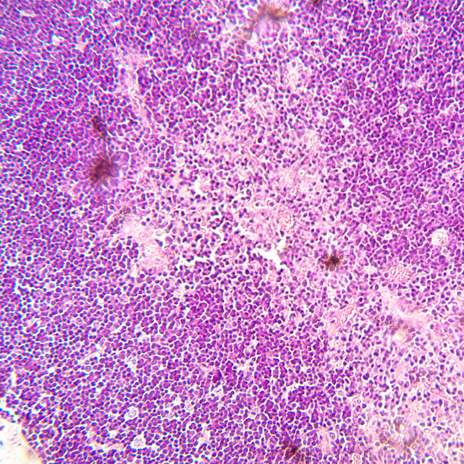 | 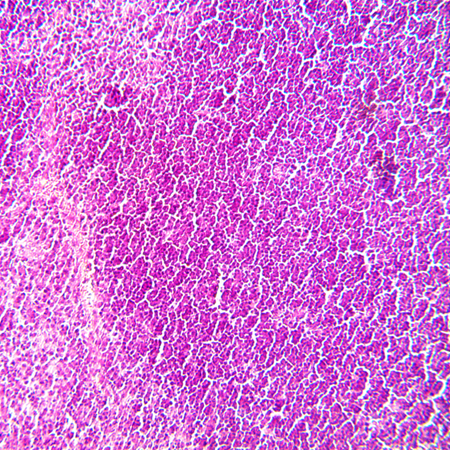 |
| Thymus, H&E, ×200 (AAV‑FST + VEGF Day 115) | Spleen, H&E, ×200 (AAV‑FST + VEGF Day 115) |
| 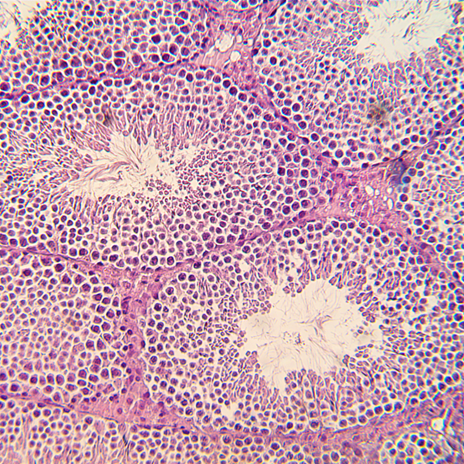 | 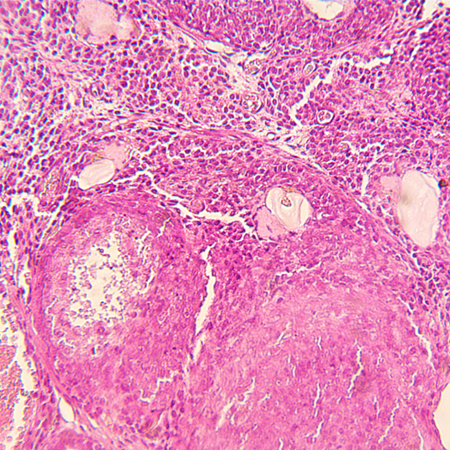 |
| Testis, H&E, ×200 (AAV‑FST + VEGF Day 115). | Ovary, H&E, ×200 (AAV‑FST + VEGF Day 115). |
| 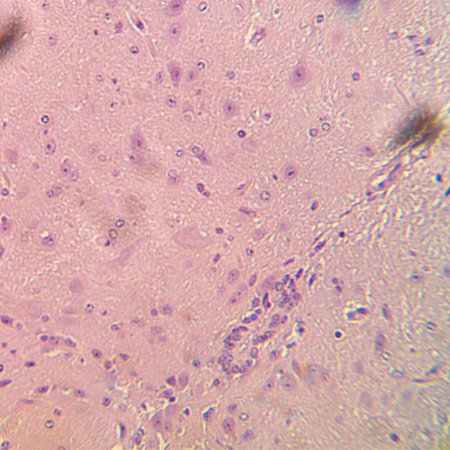 | 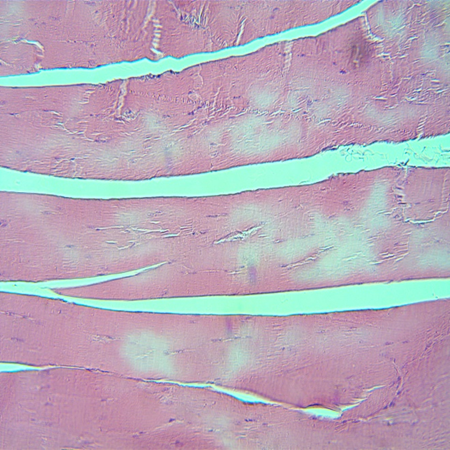 |
| Dorsal root ganglion, H&E, ×200 (AAV‑FST + VEGF Day 115) | Skeletal muscle at injection site, H&E, ×200 (AAV‑FST + VEGF Day 115). |

Representative H&E‑stained sections (×200) from the AAV‑FST (Day 25) + VEGF (Days 1 and 10) group show normal histological architecture of the heart, lung, liver, kidney, brain, thymus, spleen, gonads and dorsal root ganglia. Skeletal muscle at the injection site exhibits overall preserved morphology of cross‑striated fibres with peripheral nuclei, with focal areas of cytoplasmic clearing confined to individual fibres and no evidence of inflammatory or degenerative pathology.
